# HIV Vpr activates a nucleolar-specific ATR pathway to degrade the nucleolar stress sensor CCDC137

**DOI:** 10.1101/2024.09.11.612530

**Authors:** Karly A Nisson, Rishi S Patel, Yennifer Delgado, Mehdi Bouhaddou, Lucie Etienne, Oliver I Fregoso

## Abstract

The lentiviral accessory protein Vpr engages an extensive network of cellular pathways to drive diverse host consequences. Of its many phenotypes, CRL4A-E3 ubiquitin ligase complex co-option, DNA damage response (DDR) engagement, and G2/M arrest are conserved and thus proposed to be functionally important. How Vpr effects these functions and whether they explain how Vpr dysregulates additional cellular pathways remain unclear. Here we leverage the ability of Vpr to deplete the nucleolar protein CCDC137 to understand how Vpr-induced DDR activation impacts nucleolar processes. We characterize CCDC137 as an indirect Vpr target whose degradation does not correlate with Vpr-induced G2/M arrest. Yet, degradation is conserved among Vpr from the pandemic HIV-1 and related SIVcpz/SIVgor, and it is triggered by genomic insults that activate a nucleolar ATR pathway in a manner similar to camptothecin. We determine that Vpr causes ATR-dependent features of nucleolar stress that correlate with CCDC137 degradation, including redistribution of nucleolar proteins, altered nucleolar morphology, and repressed ribosome biogenesis. Together, this data distinguishes CCDC137 as a non-canonical Vpr target that may serve as a sensor of nucleolar disruption, and in doing so, identifies a novel role for Vpr in nucleolar stress.

**GRAPHICAL ABSTRACT:** 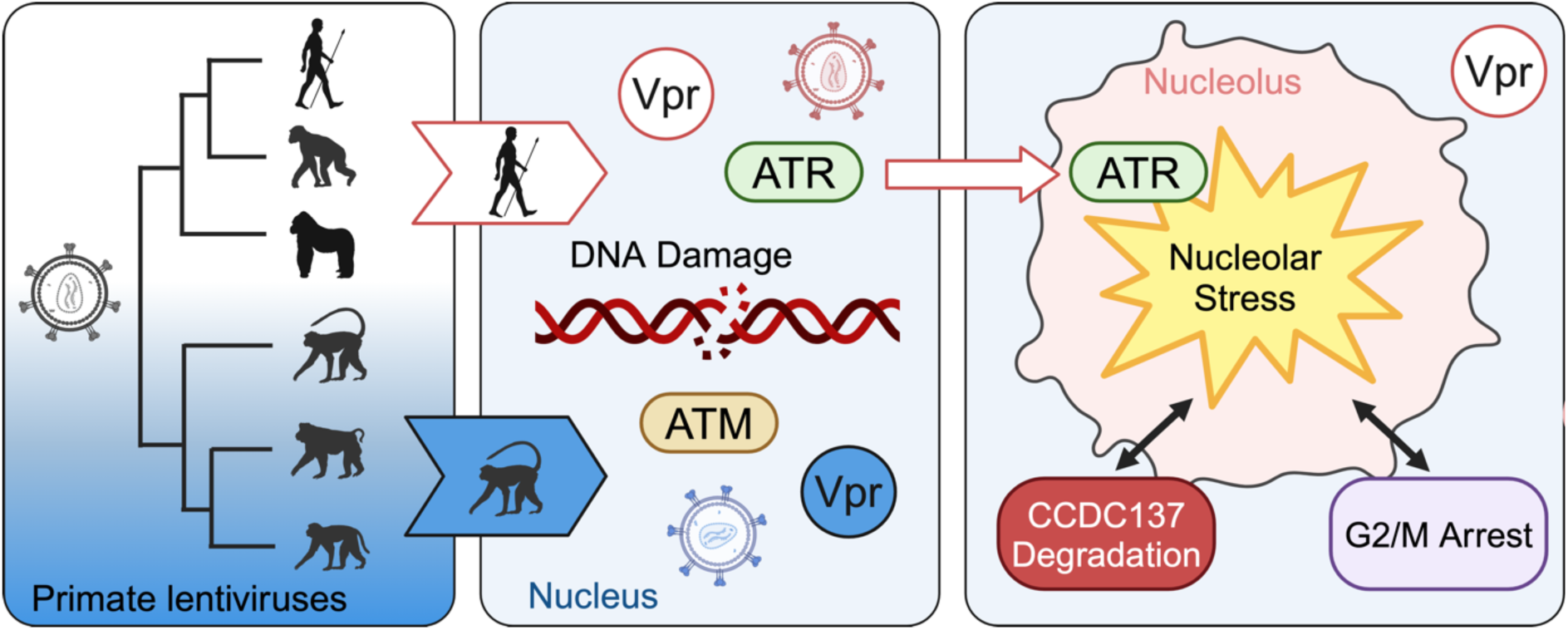

## INTRODUCTION

Lentiviral accessory proteins inhibit cellular factors and pathways that interfere with viral replication to counteract host innate immunity or augment pro-viral signaling (1). Of the four HIV-1 accessory proteins – Vpr, Vif, Vpu and Nef – Vpr is the only factor that has not yet been associated with a primary target(s) or function(s). Despite this, Vpr is evolutionarily conserved among all extant primate lentiviruses (2, 3) and essential for *in vivo* infection by HIV and SIV (4–7).

Although lentiviral Vpr has not yet been assigned a primary phenotype, the many targets and functions ascribed to it have cemented Vpr as a highly versatile protein with extensive cellular consequences. Its diverse roles include binding the host factor DCAF1 to co-opt the CRL4A E3 ubiquitin ligase complex and subsequently tag host proteins for degradation, arresting the cell cycle, promoting apoptosis, augmenting viral transcription, and engaging the DNA damage response (DDR) (8). Central to these diverse functions is the host DDR, which Vpr engages through multiple, distinct mechanisms. Of the three primary DDR signaling cascades – DNA-dependent protein kinase (DNA-PK), ataxia telangiectasia mutated (ATM), and Ataxia telangiectasia and Rad3 (ATR) kinases – Vpr activates both ATM and ATR (9–13), triggering downstream ATM-mediated NF-KB activation (14) and ATR-dependent G2/M cell cycle arrest (11), respectively. Yet, despite requiring ATR activation (11, 15), Vpr-induced cell cycle arrest cannot be explained by damage alone, as HIV-1 mutants lacking the ability to cause G2/M arrest are able to activate DDR signaling (13). In addition to activating these kinases, Vpr further engages the host DDR by causing DNA damage, inhibiting DNA repair, and binding and degrading many proteins involved in the DDR (reviewed in (16)).

CCDC137 is a recently identified Vpr target whose degradation has been connected to Vpr-induced DNA damage, G2/M arrest, and increased viral transcription (17). The precise cellular pathways that connect CCDC137 to these phenotypes, however, are unclear. Although CCDC137 functions are still emerging, current literature depicts CCDC137 as a multi-faceted nucleolar protein associated with diverse functions and processes. Multiple studies implicate CCDC137 overexpression in cancer progression and poor clinical outcomes (18–22), suggesting it plays an important role in regulating cellular homeostasis. In hepatocellular carcinoma (HCC), for instance, RNA binding (20) and E3 ubiquitin ligase co-option (19) by CCDC137 alter phosphoinositide 3-kinase (PI3K)/Akt signaling to promote cancer progression. In addition to its roles in tumorigenesis, CCDC137 modulates nuclear receptor signaling by sequestering the estrogen related (ERR) and retinoic acid (RAR) receptors in the nucleolus (23, 24). The adenoviral E1A protein binds CCDC137 to disrupt its association with RAR, thus releasing RAR from the nucleolus to engage in downstream signaling (23). Together these findings characterize CCDC137 as a versatile, though enigmatic, protein with ties to cellular proliferation. None, however, fully explain how Vpr-mediated CCDC137 degradation might contribute to Vpr-associated phenotypes, nor do they describe how CCDC137 might operate in the nucleolar compartment it inhabits.

The nucleolus is a dynamic subnuclear structure that serves as the site for the initial stages of ribosome biogenesis. Its organization is intrinsically linked to this process, encompassing three different layers representing successive steps of ribosome biogenesis, including RNA Pol I-driven ribosomal RNA (rRNA) transcription, rRNA processing, and ribosome assembly (25). Because of the cell’s reliance on this process, the nucleolus is an essential central hub for coordinating the stress response to a variety of cellular insults (26). Nucleolar perturbation by insults such as DNA damage, nucleotide depletion, oncogenic abnormalities, or viral infections (26, 27) are sufficient to rapidly trigger nucleolar stress that can lead to cell cycle arrest or apoptosis (28). In addition, more recent findings demonstrate a role for the nucleolus in orchestrating cellular stress pathways that go beyond a response related to ribosome biogenesis, including DDR regulation and maintenance of genome stability (29). In line with this, a tremendous amount of crosstalk exists between the nucleolus and DDR, as many DNA repair proteins localize to the nucleolus and, conversely, many nucleolar proteins function in the DDR (29). Both ATM and ATR have been implicated in the nucleolar response to DDR activation depending on the cellular insult (30).

Although Vpr dysregulates many of the pathways implicated in nucleolar stress, a role for Vpr in nucleolar processes has yet to be defined. Here we leverage the connections between CCDC137, the nucleolus, and the DDR to test the hypothesis that Vpr modulates nucleolar pathways. We find that CCDC137 is an indirect Vpr target whose degradation is associated with, but not a direct cause or consequence of, Vpr-induced G2/M arrest. Additionally, we characterize CCDC137 as an evolutionarily important Vpr target that bridges multiple Vpr phenotypes, including a newly identified Vpr function, nucleolar stress. We further distinguish CCDC137 as a nucleolar stress sensor that is degraded when Vpr-induced DNA damage triggers the activation of a nucleolar-specific ATR pathway. Together, these results characterize the evolutionary importance of CCDC137 depletion to Vpr function, define DDR activation as required for this depletion, and identify a role for Vpr in nucleolar processes.

## MATERIAL AND METHODS

### Plasmids and generation of viruses

pcDNA3.1-3XFLAG-Vpr, pcDNA3.1-3XFLAG-Vpx, pscAAV-GFP-T2A-Vpr, pscAAV-mCherry-T2A-Vpr, and LPCX-HA-SAMHD1 plasmids were described previously (2, 31). Vpr and Vpx sequences represent the following isolates: HIV-1 Bru (H1B.FR.83.HXB2_LAI_IIIB_BRU; GenBank K03455.1), HIV-1 Q23-17 (H1.A.Q2317; GenBank AF004885.1), HIV-1 ETH220 (H1C.ET.86.ETH2220; GenBank U46016.1), HIV-1 SE6165 (H1G.SE.93.SE6165 GenBank: AF061642.1), HIV-1 SE9280 (H1J.SE.93.SE9280; GenBank AF082394.1), HIV-1 P Fr (P RBF168; GenBank GU111555.1), HIV-2 Rod9 (H2A.Rod9; GenBank M15390.1), HIV-2 7312a (H2.B.7312a; GenBank L36874.1), HIV-2 A.PT.x.ALI (H2A.PT.x.ALI; GenBank AF082339.1); HIV-2 AB.JP.04 (H2_01_AB.JP.04.NMC307_20; GenBank AB731738.1), HIV-2 B.GH.86 (H2B.GH.86.D205; GenBank X61240.1), HIV-2 ABT96 (H2G.Cl.92.ABT96; GenBank AF208027.1), SIVcpz MB897 (CPZ.CM.x.SIVcpzMB897; GenBank EF535994.1), SIVcpz EK505 (CPZ.CM.05.SIVcpzEK505; GenBank DQ373065.1), SIVcpz CAM13 (CPZ.CM.01.SIVcpzCAM13; GenBank AY169968.1), SIVgor CP684 (GOR.CM.04.SIVgorCP684com.; GenBank FJ424871), SIVsmm SL9 (SMM.SL.92.SL92B; GenBank AF334679.1), SIVsmm CFU212 (SMM.US.86.CFU212; GenBank JX860407.1), and SIVmac 239 (MAC.US.x.239; GenBank M33262). In addition, the following consensus Vpr sequences were used: HIV-1 N con (32), HIV-1 O con (32), and HIV-1 P con (MEQAPEDQGPPREPFNEWMLQTLEELKAEAVRHFPMPWLHSLGQYIYNTYGDTWEGVTAIIRILQQLIFIHYRIGCQHSRIGILTLSRRGGRHGPSRS).

LPCX-HA-CCDC137 plasmids were generated by synthesizing CCDC137 as gBlocks (IDT) that were subcloned into LPCX-SAMHD1-HA (2) using standard cloning techniques. CCDC137 protein sequences are available at Figshare: https://doi.org/10.6084/m9.figshare.26841640 and https://doi.org/10.6084/m9.figshare.26841634.

pscAAV-mCherry-T2A Vpr and pscAAV-GFP-T2A Vpr plasmids were packaged into rAAV vectors through transient transfections of HEK 293 cells using polyethyleneimine (PEI) as previously described (33). Inverted terminal repeat (ITR)-specific quantitative PCR (qPCR) was used to quantify levels of DNase-resistant vector genomes as previously described (34).

### Cell lines and cell culture

Human embryonic kidney (HEK) 293 and human bone osteosarcoma epithelial (U2OS) cells were cultured as adherent cells in Dulbecco’s modified Eagle’s medium (DMEM) growth medium (high glucose, L-glutamine, no sodium pyruvate) with 10% fetal bovine serum (Peak Serum) and 1% penicillin-streptomycin (Gibco) at 37°C and 5% CO_2_. All cells were harvested using 0.05% trypsin-EDTA (Gibco). Transfections were performed with TransIT-LT1 (Mirus), and rAAV infections were performed by adding rAAV directly to the growth medium. For inhibitor assays, cells were incubated with 5nm Actinomycin D (Gibco) for 5 hours (hr), 5µM or 10µM MLN4924 (Cell Signalling) for 8hr, 10µM MG132 (Sigma) for 5hr, or 10µM ATMi (KU-55933) (Cayman Chemicals) or ATRi (VE-821) (Cayman Chemicals) for 24hr. For genotoxic agent assays, cells were incubated with titrating amounts of camptothecin (Cayman Chemicals), cisplatin (Tokyo Chemical Industry), etoposide (Cayman Chemicals), hydroxyurea (Sigma) or Olaparib (MedChemExpress) for 24hr.

### Flow Cytometry

Approximately 6.0 x 10^6^ cells per condition were harvested at 24 hours post infection (hpi) and stained with Hoescht 33342 Ready Flow^TM^ Reagent (Invitrogen) according to the manufacturer’s instructions for cell cycle analysis. Event counts were measured on the AttuneNxT (Thermo Fisher Scientific), and data were analyzed using FlowJo software. G1 and G2 values were obtained using the Dean-Jett-Fox Univariate model (35, 36), and G2/G1 ratios were calculated from these percentages.

### Western blots and co-immunoprecipitations

Cells were lysed for western blot in radioimmunoprecipitation assay (RIPA) buffer (50mM Tris-HCl [pH 8.0], 150mM NaCl, 1mM EDTA, 0.1% SDS, 1% NP-40, 0.5% sodium deoxycholate, Universal Nuclease (Pierce), and EDTA-free protease inhibitor (Pierce) and clarified by centrifugation at 15,000 x *g* for 15 min. Immunoprecipitations were performed as previously described (32) using anti-HA affinity gel (Millipore Sigma) in NP-40 lysis buffer (200mM NaCl, 50mM Tris [pH 7.4], 0.5% NP-40, 1mM DTT, Universal nuclease, protease inhibitor). All samples were boiled in 4X sample buffer (40% glycerol, 240mM Tris, pH 6.8, 8% SDS, 0.5% β-mercaptoethanol, and bromophenol blue), run on 4 to 12% Bis-Tris polyacrylamide gels, and subsequently transferred onto polyvinylidene difluoride membranes (Sigma). Membranes were blocked in intercept blocking buffer (Li-COR Biosciences) and probed with the indicated primary antibodies: mouse anti-FLAG M2 1:3000 (Sigma), mouse anti-HA 1:10000 (Invitrogen), rabbit anti-CC137 1:2000 (Atlas and Abcam), rabbit anti-VprBP 1:2000 (Proteintech), mouse anti-β-tubulin 1:5000 (Cell Signaling), rabbit anti-γH2AX 1:5000 (Cell Signaling), and rabbit anti-p53 1:5000 (Proteintech). Membranes were incubated with secondary antibodies IRDye 800CW anti-Rabbit and IRDye 680RD anti-mouse 1:20000 (Li-COR Biosciences) and visualized using the Li-COR Odyssey M (Li-COR Biosciences).

#### Quantification of CCDC137 protein levels

Degradation experiments were performed as biological triplicates (*n=3*), and protein levels were quantified using ImageStudio (Li-COR Biosciences). Each condition was normalized to a corresponding β-tubulin loading control. CCDC137 degradation values for a given Vpr are reported as the percentage of CCDC137 protein remaining relative to an empty vector control, averaged across three separate experiments. From these percentages, each degradation phenotype is then classified as robust (0-35% CCDC137 remaining), intermediate (36-80% CCDC137 remaining) or absent (81-100% CCDC137 remaining); these descriptions are used throughout the text to represent calculated values that fall within these ranges. Degradation heatmaps were generated based on the averaged percentages from three biological replicates (*n=3*).

### Quantitative Reverse Transcription PCR (qRT-PCR)

Cells were harvested at 24hpi and RNA was isolated using the PureLink^TM^ RNA Mini Kit (Invitrogen). cDNA was generated using the SuperScript IV First-Stand Synthesis System (Invitrogen) with Oligo(dT) primers. qPCR reactions were prepared using PowerTrack SYBR Green Master Mix (Thermo Fisher Scientific) and the following primers (5’ to 3’): CCDC137 TATGAGGAGCCGCCAAGAGATG and CCTCTCCCTTTGCTTCCTTCTC, and GAPDH CAAGATCATCAGCAATGCCT and AGGGATGATGTTCTGGAGAG (IDT). All reactions were carried out on the LightCycler 480 System (Roche). To quantify mRNA levels, ΔΔCt values were calculated for each sample, and CCDC137 values were normalized to ΔΔCt values for the housekeeping gene GAPDH. Statistical data were based the averaged normalized ΔΔCt values from triplicate reactions performed for three different biological replicates.

### Immunofluorescence

Cells were plated onto glass slides in 24-well plates (Greiner Bio-One) at 1.5×10^5^ cells per well and allowed to expand overnight. For localization and nucleolar size assays, cells were harvested at 24hr post-treatment. Cells were permeabilized (0.5% Triton-X 100 in PBS) for 5min on ice prior to fixation (4% paraformaldehyde in PBS) for 20min at 20°C and then incubated overnight in PBS at 4°C. Cells infected with the rAAV HIV-1 Q65R mutant were fixed prior to permeabilization. Coverslips were then washed three times in PBS before blocking (30% BSA, 0.1% Tween 20 in PBS) for 30min at 20°C. Cells were incubated with the following primary antibodies diluted in blocking buffer overnight at 4°C: rabbit anti-CC137 1:200 (Abcam), rabbit anti-NPM1 1:500 (Atlas), rabbit anti-Nucleolin 1:1000 (Cell Signaling) and mouse anti-FLAG M2 1:5000 (Sigma). Coverslips were then washed three times in 0.1% Tween 20 in PBS for 5min each at 20°C and incubated with Alexa Fluor-conjugated secondary antibodies (Life Technologies) for 1hr at 20°C. Nuclei were stained with Hoescht 33342 1:2000 (Invitrogen) diluted in PBS. Coverslips were mounted onto glass slides using antifade mounting media (Invitrogen).

For 5-ethynyl uridine (EU) experiments, cells were harvested at 28 hpi or 24hr post drug treatment following a 1hr incubation with 0.8mM EU (Invitrogen). Coverslips were prepared according to manufacturer’s instructions for the Click-iT™ RNA Alexa Fluor™ 488 Imaging Kit (Invitrogen).

Representative images for all IF experiments were acquired on the LSM 980 using Airyscan. For 5-EU and nucleolar size assays, images for quantification were obtained using the Zeiss Axio Imager Z1. Fiji was used to quantify nucleolar area and EU signal, measured as mean fluorescence intensity (MFI) of total nuclei as previously described (37). Statistical analyses were based on approximately 100-200 cells per condition.

### Phylogenetics and positive selection (PS) analyses

Phylogenetics, positive selection analysis, and detection of other potential genetic innovations (such as recombination, insertions/deletions and duplications) were performed using the Detections of Genetic Innovations (DGINN) pipeline, as previously described (38). Briefly, homologous codon sequences to human CCDC137 in primates were automatically retrieved using NCBI blastn implemented in DGINN. Sequences were aligned using Prank (39) and recombination events were detected using Genetic Algorithm for Recombination Detection (GARD) (40). Amino acid alignment at the primate level was also generated using Muscle (41). Positive selection marks were detected in a simian codon Prank alignment of CCDC137 orthologs from 23 species using the following tests of selection in DGINN: HYPHY BUSTED and MEME, PAML codeml (M0, M1, M2, M7, M8, M8a), and Bio++ (M0, M1^NS^, M2^NS^, M7^NS^, M8^NS^, M8a^NS^)(42–45). Sites under positive selection were identified using HYPHY MEME (*P* value < 0.05), from the codeml M2 and M8 model BEB statistics (BEB > 0.95), and Bio++ Bayesian posterior probabilities (PP) from the M2^NS^ and M8^NS^ models (PP > 0.95). In addition, Fast, Unconstrained Bayesian AppRoximation for Inferring Selection (FUBAR) web-based method was also used to detect site-specific positive selection (PP > 0.90) (46). Alignments are openly available at FigShare: https://doi.org/10.6084/m9.figshare.26841640 and https://doi.org/10.6084/m9.figshare.26841634.

### Protein structure prediction with AlphaFold

The AlphaFold predicted structure of the human CCDC137 protein (accession Q6PK04) was obtained from the AlphaFold database (https://alphafold.ebi.ac.uk/entry/Q6PK04) as no experimentally determined structures were available. ChimeraX was used to annotate the default CCDC137 predicted structure.

### Proteomic Analyses

For gene set over-representation analysis (GSOA), genes from the Greenwood *et al.* dataset (47) or the Johnson *et al.* dataset (48) were selected if they possessed a log2 fold change of at least 0.5 in either direction. GSOA was performed using the enricher function of clusterProfiler package (version 3.12.0) in R with default parameters. Significant gene ontology (GO) terms (adjusted p value < 0.05) were identified and further refined to select non-redundant terms. To select non-redundant gene sets, a GO term tree was first constructed based on distances (1-Jaccard Similarity Coefficients of shared genes in MSigDB) between the significant terms. The GO term tree was cut at h = 0.2 to identify clusters of non-redundant gene sets. For results with multiple significant terms belonging to the same cluster, the most significant term was selected (i.e., lowest adjusted p value) with the largest number of genes (i.e., to select the broadest categories). For the gene set enrichment analysis (GSEA), non-thresholded log2 fold changes from the Johnson *et. al* dataset were used as inputs. GO terms were not further refined. GO terms were obtained from the c5 category of Molecular Signature Database (Human MSigDB v2024.1.Hs) (49). Cytoscape (50) was used to visualize CCDC137 interactions determined using STRING (51).

## RESULTS

### CCDC137 degradation is neither a direct cause nor consequence of G2/M arrest

To investigate the connection between CCDC137 degradation and Vpr-induced G2/M arrest, we first tested whether HIV-1 Vpr mutants that lack the ability to cause cell cycle arrest could still degrade CCDC137. These mutants are made in the backbone of the primary HIV-1 isolate Q23-17 (WT) and crucially separate conserved Vpr phenotypes. Consequently, they allow us to correlate CCDC137 degradation with individual Vpr functions, including G2/M arrest (H71R, S79A, R80A) and decreased DCAF1 binding (H71R) (14). As a negative Vpr control, we also assayed the degradation capacity of the functionally dead Q65R mutant, which does not cause arrest, bind DCAF1, or properly localize within the nucleus (14). To test this, we delivered either HIV-1 WT Vpr or mutants to U2OS cells via a recombinant adeno-associated virus (rAAV) vector system (13) and assayed Vpr-induced cell cycle arrest via flow cytometry (**Figure 1A**) and endogenous CCDC137 levels via quantitative western blot (**Figure 1B-C**).

**Figure 1.**
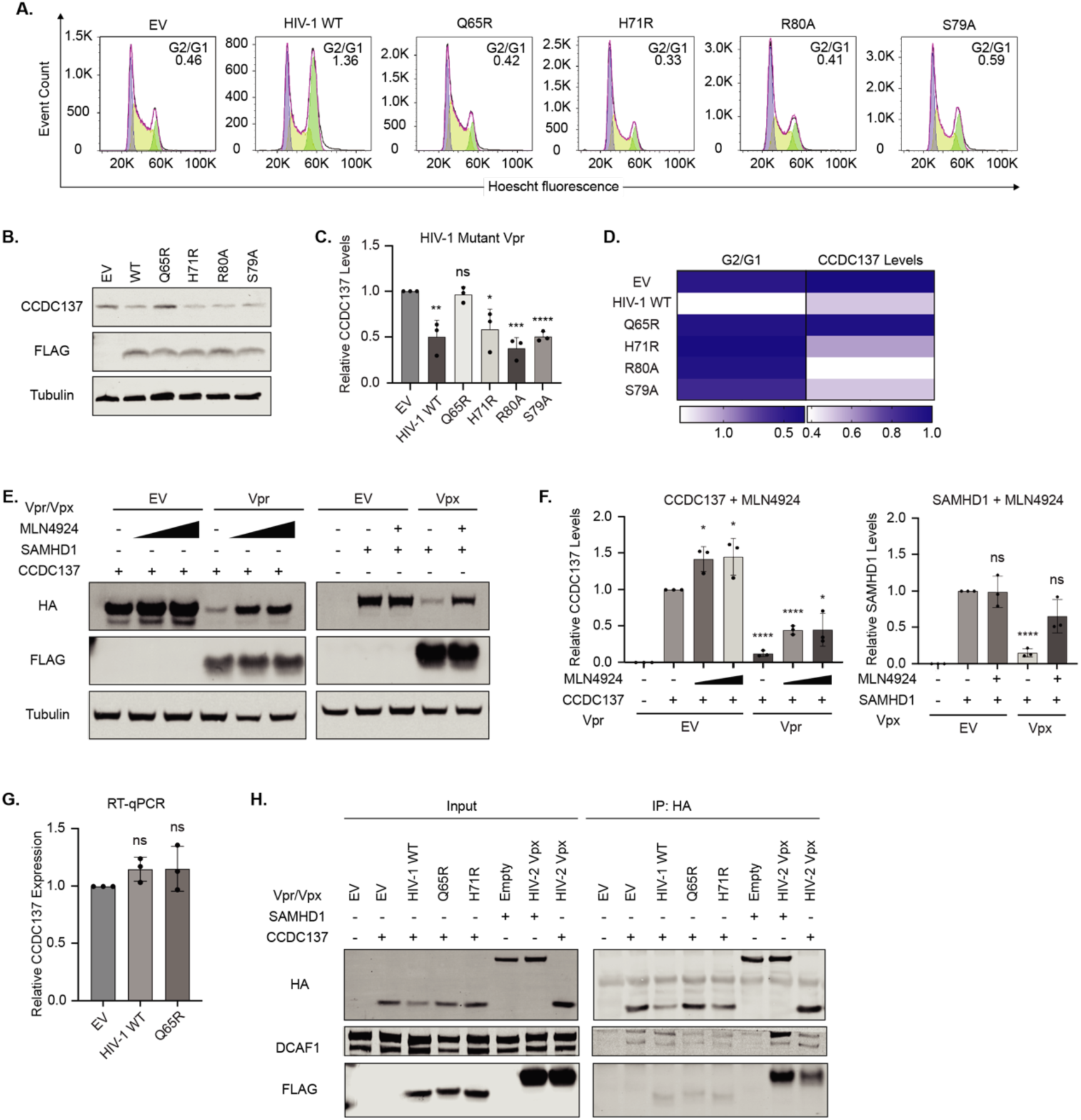
CCDC137 degradation does not correlate with Vpr-mediated G2/M arrest or CRL4A^DCAF1^ E3 ubiquitin ligase complex recruitment. (**A**) Representative flow cytometry cell cycle plots of U2OS cells infected with the indicated WT or mutant Vpr and gated for Vpr (mCherry) expression at 28 hours post infection (hpi). G2/G1 values were calculated from the percentages of cells in G2 and G1 in each condition based on DNA content (Hoechst fluorescence) (*n=3*). (**B**) Representative western blot of cells harvested from the same experiment as in (A) probed for endogenous CCDC137 and 3XFLAG Vpr. (**C**) Quantification of CCDC137 protein levels from (B) normalized to Tubulin, and relative to the empty vector control. Data from 3 biological replicates are shown. (**D**) Heatmap of G2/G1 values from (A) and average CCDC137 levels from (B) in the presence of the indicated Vpr. Color gradients were set to empty vector values (dark blue) and either the lowest CCDC137 level or highest G2/M value (white). (**E**) MLN4924 rescue of Vpr-mediated CCDC137 degradation or Vpx-mediated SAMHD1 degradation in U2OS cells treated with MLN4924 (10µM for SAMHD1 and 5µM or 10µM for CCDC137, as indicated). Western blot is representative of three separate experiments (*n=3*) and probed for HA-CCDC137 or HA-SAMHD1, 3XFLAG Vpr or Vpx, and Tubulin. (**F**) Quantification of (E) as described for (C). (**G**) qRT-PCR of endogenous CCDC137 mRNA levels in the presence of HIV-1 WT or Q65R Vpr. (**H**) Co-immunoprecipitation assays against HA-SAMHD1 or HA-CCDC137 in U2OS cells treated with 10µM MLN4924, probed for HA, 3X FLAG Vpr or Vpx, and Tubulin. All results were analyzed by unpaired t-tests. Asterisks indicate statistical significance from empty vector control; error bars represent ± standard deviation (ns = not significant, *P ≤ 0.0332, **P ≤ 0.0021, *** P ≤ 0.0002, ****P < 0.0001). EV=empty vector.

In the presence of HIV-1 WT Vpr, we observed a correlation between G2/M arrest and CCDC137 degradation as previously reported (17, 52). HIV-1 WT Vpr degraded endogenous CCDC137 (50%) and resulted in potent cell cycle arrest (G2/G1 = 1.36) when compared to an empty vector control (G2/G1 = 0.46) (**Figure 1A-C**). The functionally dead HIV-1 Q65R Vpr mutant failed to either cause arrest (G2/G1 = 0.42) or degrade CCDC137 (96%), as expected. When we introduced other HIV-1 Vpr mutants, however, we saw that all significantly degraded CCDC137 despite their inability to arrest cells in G2/M. The cell cycle arrest-deficient mutants HIV-1 R80A, S79A, and H71R resulted in arrest profiles similar to the empty vector control (G2/G1 = 0.41, 0.59, and 0.33, respectively) but degraded CCDC137 to levels induced by HIV-1 WT Vpr (38%, 51%, and 66%, respectively). Comparing the degradation and G2/G1 values side-by-side in a heatmap further highlighted a lack of correlation among the two phenotypes (**Figure 1D**). Together, this lack of correlation shows that Vpr-mediated CCDC137 degradation is neither a direct cause nor consequence of cell cycle arrest.

### CCDC137 degradation is not mediated by canonical Vpr engagement of the CRL4-E3 ubiquitin ligase complex

We next asked whether recruitment of the CRL4A^DCAF1^ E3 ubiquitin ligase complex was required for CCDC137 degradation, consistent with other canonical Vpr targets (53). First, we assayed whether this degradation was cullin-ring ligase (CRL) dependent by co-expressing hemagglutinin (HA)-tagged human CCDC137 and 3XFLAG-tagged Vpr in the presence or absence of increasing concentrations of the neddylation inhibitor MLN4924. As a positive control, we assayed MLN4924 rescue of Vpx-mediated SAMHD1 degradation, which is also CRL4A^DCAF1^-dependent. MLN4924 increased CCDC137 levels in the presence of Vpr by 3.75-fold (12% to 45% increase), similar to the positive control SAMHD1 (4.5-fold, from 15% to 66%) (**Figure 1E-F**). However, unlike SAMHD1, MLN4924 increased CCDC137 levels in cells lacking Vpr (1.45-fold change, from 100% to 145%), suggesting that basal CCD137 levels may be regulated by a cullin-dependent pathway independent of Vpr. Vpr-depleted CCDC137 levels were also rescued by the proteosome inhibitor MG132 (**Figure S1A-B**), and CCDC137 RNA levels were not increased in Vpr-treated samples (**Figure 1G**). Together, this suggests that CCDC137 levels are regulated post-transcriptionally in a cullin-dependent manner, both dependent and independent of Vpr.

To further clarify a role for CRL4A^DCAF1^ ubiquitin ligase complex recruitment in Vpr-mediated CCDC137 degradation, we next performed co-immunoprecipitations (co-IPs) to probe for the recruitment of DCAF1, the adaptor protein between Vpr and CRL4A-DDB1 (54), to a Vpr-CCDC137 complex. In the presence of MLN4924, we transiently co-expressed HA-tagged human CCDC137 and 3X-FLAG tagged Vpr in U2OS cells, performed IPs against HA-CCDC137, and probed for interactions with 3X-FLAG Vpr and endogenous DCAF1. As has been previously reported, HIV-2 Rod9 Vpx robustly interacted with human SAMHD1, and DCAF1 was only recruited in the presence of HIV-2 Rod9 Vpx (55, 56). In contrast, we only detected minimal interaction between CCDC137 and HIV-1 WT Vpr. The intensity of this interaction was no greater than that observed between CCDC137 and HIV-2 Rod9 Vpx, which does not degrade CCDC137 (**Figure 1H**). Moreover, low levels of DCAF1 were already bound to CCDC137 in the absence of Vpr when compared to an empty vector control, and these levels did not increase in the presence of HIV-1 WT Vpr. HIV-1 Vpr mutants that could or could not degrade CCDC137, H71R and Q65R, respectively, showed similar weak band intensities for 3XFLAG-Vpr and DCAF1 that we observed with WT Vpr. Consistent with this minimal interaction, we also did not observe co-localization among Vpr and CCDC137 via immunofluorescence (IF), despite dimmer nucleolar CCDC137 signal in cells expressing Vpr orthologs that were capable of degradation (**Figure S1C**).

Overall, Vpr depleted CCDC137 in the absence of a robust direct interaction, additional DCAF1 recruitment, or altered CCDC137 localization. While cellular CCDC137 levels may be regulated via a DCAF1-dependent mechanism, Vpr is not enhancing this interaction. Together, this suggests that Vpr does not degrade CCDC137 via canonical co-option of the CRL4A^DCAF1^ ubiquitin ligase complex.

### The ability to deplete CCDC137 is evolutionarily conserved among Vpr orthologs from the HIV-1 lineage

Although we did not find evidence for a direct correlation between Vpr-mediated CCDC137 degradation and G2/M arrest, we reasoned that this degradation may still be an evolutionarily conserved, and thus important, capacity of Vpr. We first asked whether CCDC137 degradation was conserved within the broader Vpr diversity from HIV-1 and HIV-2, which both infect humans but have distinct evolutionary origins: HIV-1 originating from spillovers of SIVcpz and SIVgor, and HIV-2 from SIVsmm (57). Using transient co-transfections of HA-CCDC137 and 3X-FLAG Vpr, we determined that four Vpr isolates from the pandemic HIV-1 group M viruses robustly degraded human CCDC137 (all < 20%), as did a consensus Vpr from group O (3.5%). A consensus Vpr from group N and two Vpr isolates from group P (one a primary sequence and one a consensus Vpr) did not significantly degrade CCDC137 (68%, and 96 and >100%, respectively) (**Figure 2A-B**). In contrast to HIV-1 Vpr, HIV-2 Vpr isolates showed intermediate or no degradation of human CCDC137, while Vpx showed no degradation (>35% CCDC137 remaining) (**Figure S2A-B**), suggesting this function is not present among HIV-2 viruses.

**Figure 2.**
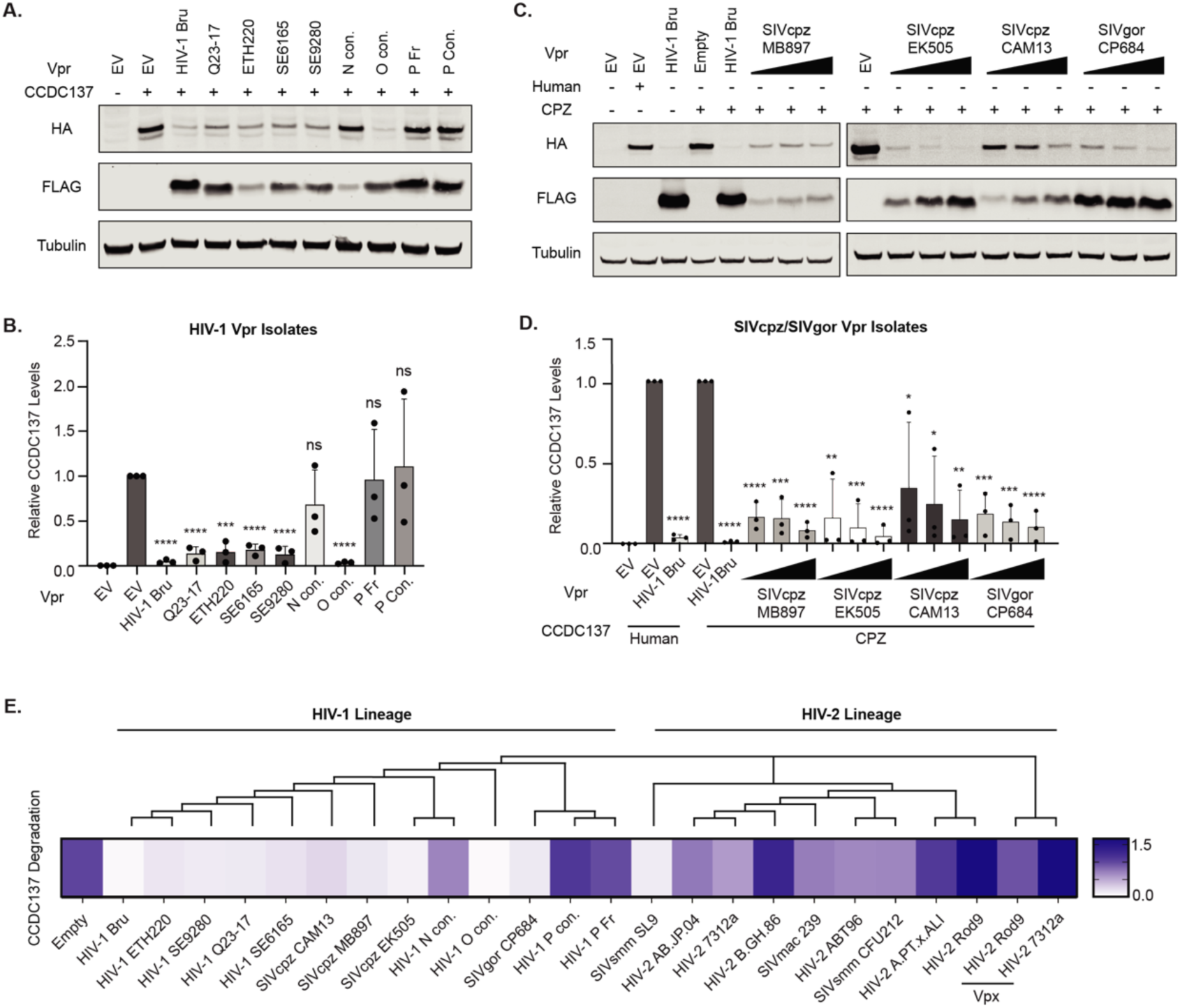
Degradation of CCDC137 is conserved among the HIV-1/SIVcpz/SIVgor Vpr lineage. (**A**) Representative western blots of U2OS cells transiently transfected with human HA-CCDC137 and 3XFLAG HIV-1 Vpr isolates. Cells were harvested at 28 hours post-transfection, and western blots were probed and (**B**) quantified as in Figure 1C (*n=3*). (**C**) Representative western blots and (**D**) quantification of chimpanzee (CPZ) HA-CCDC137 degradation by 3XFLAG SIVcpz or SIVgor Vpr isolates as described in (A) and (B) (*n=3*). (**E**) Heatmap of HA-CCDC137 levels in the presence of Vpr/Vpx from the HIV-1 or HIV-2 lineage. The color gradient was set to the highest (dark blue) and lowest (white) CCDC137 levels measured. The corresponding cladogram represents the aligned sequences of the Vpr/Vpx orthologs used in (A-D) and SF2 (A-D) as indicated. Results were analyzed by unpaired t-tests. Asterisks indicate statistical significance from empty vector control; error bars represent ± standard deviation (ns = not significant, *P≤0.0332, **P≤0.0021, *** P≤0.0002, ****P<0.0001). EV = empty vector; con. = consensus.

We next explored whether the ability of Vpr to degrade CCDC137 was conserved among the ancestral lineages of HIV-1 and HIV-2 viruses. Within the HIV-1 lineage, we found that all Vpr isolates tested from SIV infecting chimpanzees (SIVcpz) and gorilla (SIVgor) robustly degraded chimpanzee (*Pan troglodytes troglodytes*) CCDC137 at each concentration tested (**Figure 2C-D**). Among the HIV-2 lineage, SIV sooty mangabeys (SIVsmm) Vpr isolate SL9, which is phylogenetically distant from the other HIV-2 and SIVsmm isolates, robustly degraded CCDC137 from sooty mangabeys (*Cercocebus atys*). However, SIVsmm CFU212 and SIVmac239 (from macaque) Vpr had intermediate degradation phenotypes, in line with other HIV-2 isolates (**Figure S2C-D**). These results suggest that the ability to degrade CCDC137 is conserved among Vpr from HIV-1/SIVcpz/gor but not the HIV-2 lineage and may thus be a specific and important function of HIV-1 and SIVcpz/gor (**Figure 2E**).

### Specificity of the Vpr-mediated CCDC137 degradation is conferred by both the Vpr and CCDC137 sequence

To determine whether the degradation capacity of a given Vpr was inherent to the Vpr or dependent on the CCDC137, we challenged a panel of Vpr orthologs and isolates that could or could not degrade CCDC137 against the natural diversity of CCDC137 present in primates. We selected four Vpr orthologs based on their diverse abilities to degrade their autologous CCDC137 (**Figure 2**): HIV-1 Bru (robust), HIV-2 7312a (intermediate), SIVcpz MB897 (robust), and SIVsmm CFU212 (intermediate). We assayed degradation against the breadth of CCDC137 sequence variability from humans, chimpanzees, sooty mangabeys, African Green Monkeys (AGMs) (*Chlorocebus sabaeus*), a prosimian (PRO, *Propithecus coquereli*), and a New World monkey (NWM, *Callithrix jacchus*). We found that HIV-1 and SIVcpz Vpr robustly degrade nearly all CCDC137 orthologs tested, including those from prosimians and NWM. However, degradation capacities of HIV-2 7312a and SIVsmm CFU212 Vpr varied, depending on the CCDC137 ortholog tested (**Figure 3A-D and S2E-F**). Together, these data suggest that degradation is an innate capacity of certain Vpr orthologs that is also influenced by unique properties of certain CCDC137 proteins (**Figure 3E**). Although these phenotypes are not reminiscent of typical virus-host molecular arms-races, in which we might expect robust degradation phenotypes in the context of autologous hosts, there is some specificity conferred by both the CCDC137 (host) and Vpr (viral) sequence.

**Figure 3.**
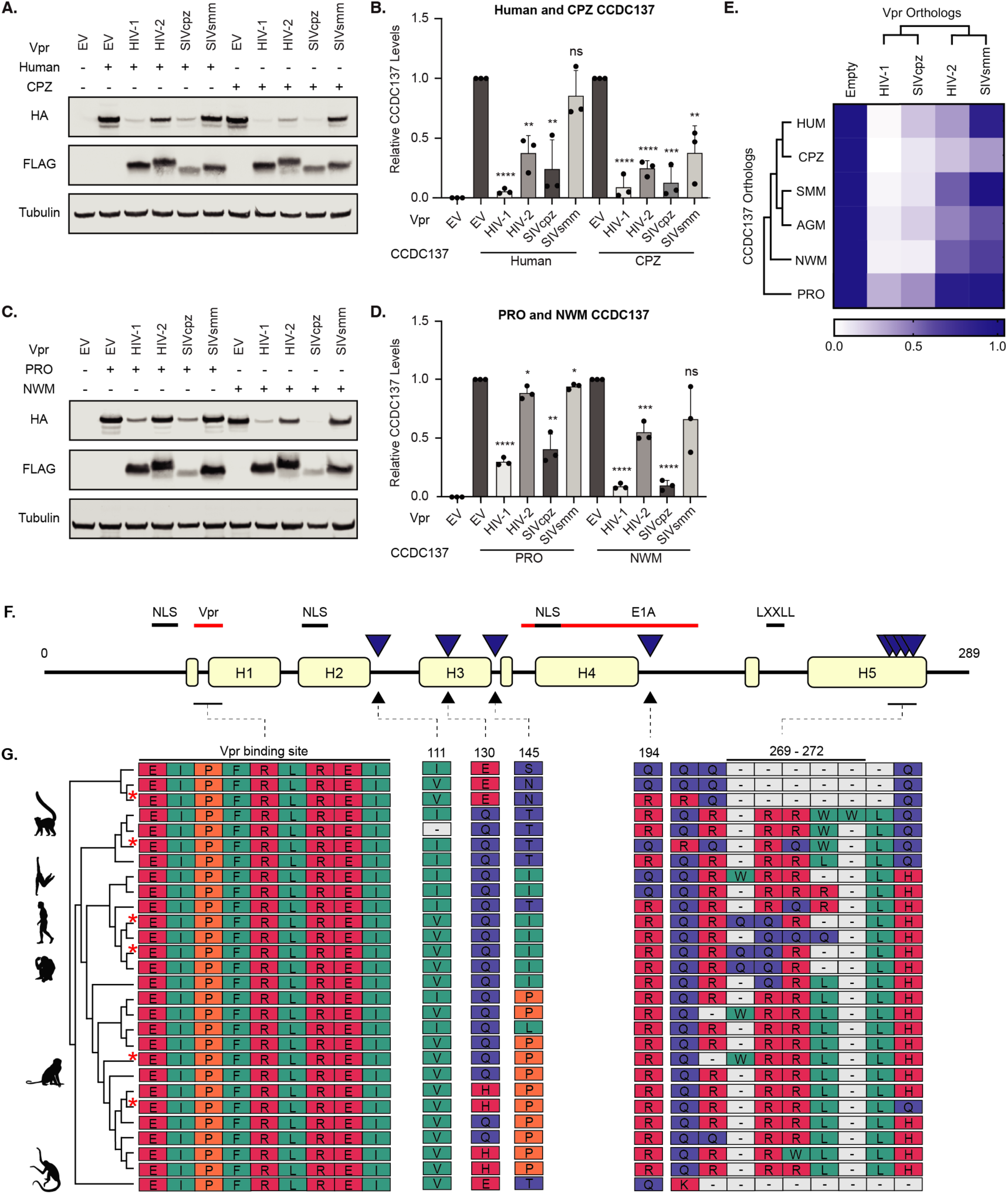
Host species- and viral lineage-specific determinants influence CCDC137 degradation. (**A**) Representative western blot and (**B**) quantification of human and chimpanzee (CPZ) HA-CCDC137 degradation by a panel of 3XFLAG HIV-1 Bru, HIV-2 7312a, SIVcpz MB897 and SIVsmm CFU212 Vpr orthologs. Assays were performed as described in Figure 2 (A-D) (*n=3*). (**C**) Representative western blot and quantification of (**D**) prosimian (PRO) and New World monkey (NWM) HA-CCDC137 degradation by the indicated Vpr (*n=3*). (**E**) Heatmap depicting degradation of the different CCDC137 orthologs (rows) by the indicated Vpr ortholog (columns). Results were analyzed by unpaired t-tests. Asterisks indicate statistical significance from empty vector control; error bars represent ± standard deviation (ns = not significant, *P≤0.0332, **P≤0.0021, *** P≤0.0002, ****P<0.0001). (**F**) Schematic representation of the CCDC137 protein. Helices are noted as H1-5 and sites of PS (Table S1 for details and methods) are represented as solid blue triangles. CCDC137 sequence motifs and viral protein binding sites are represented as labeled black and red lines, respectively. (**G**) Residues corresponding to the proposed Vpr binding site and marks of PS. Amino acids are labeled by property (pink = charged, teal = hydrophobic uncharged, blue = polar uncharged, and orange = special case). Grey boxes containing (-) indicate a lack of corresponding amino acid in the alignment. Sequences are organized based on an evolutionary tree (left) constructed from 28 primate CCDC137 sequences. Red asterisks indicate sequences used in (A-E). Primate icons are from Adobe Stock Images. EV = empty vector. NLS = nuclear localization sequence.

We therefore characterized the evolutionary history of CCDC137 in primates and tested whether it has experienced episodes of rapid evolution and positive selection (PS). We performed PS analyses using the Detection of Genetic INNovations (DGINN) pipeline, which performs phylogenetic and positive selection analyses from five complementary methods to increase the sensitivity and specificity in the detection of genetic innovation. Among simians, we did not find evidence of strong PS in CCDC137 (**File S1**). However, performing detailed site-specific analyses, we identified five regions undergoing rapid evolution (**Figure 3F-G**). Most are in predicted disordered regions, except a 269-272 region in the C-terminal helix of CCDC137 (**Figure S2G**). Furthermore, site 194 overlapped with a region linked to adenoviral E1A interaction (23), which could be reminiscent of a past evolutionary conflict. However, no signatures of positive selection or rapid evolution overlapped with nuclear localization sequences (NLS), a nuclear-hormone receptor binding motif (LXXLL), or a sequence associated with Vpr-mediated CCDC137 degradation (17). Remarkably, the latter was entirely conserved amongst primate sequences (**Figure 3F-G**). Given the functional host-specificity observed, it is thus likely that other CCDC137 determinants beyond sequence identity are also involved in CCDC137-induced degradation by Vpr.

### CCDC137 depletion is a consequence of specific genomic insults

Despite our findings that CCDC137 is not a canonical target of Vpr, it is clear that Vpr orthologs from the HIV-1 lineage potently deplete this host factor. Given the absence of a direct CCDC137-Vpr interaction, we reasoned that CCDC137 depletion might result from another important HIV-1 Vpr function. Considering that CCDC137 degradation has been linked to activation of the DDR (17) and Vpr both induces DNA damage and modulates DDR signaling, we asked whether CCDC137 degradation is a consequence of DNA damage.

To test this, we investigated whether a panel of different genotoxic agents were sufficient to deplete CCDC137. Within this panel, we assayed CCDC137 levels in the context of γH2AX levels, which we used to measure the amount of global DDR activation induced by each agent. We found that titrating amounts of camptothecin (**Figure 4A-B**) and etoposide (**Figure S3A-B**), which cause DNA damage by inhibiting Topoisomerase I and II, respectively, robustly depleted CCDC137. In contrast, cisplatin (**Figure 4C-D**), an agent that forms intra-strand DNA crosslinks that inhibit DNA synthesis, and hydroxyurea (**Figure S3C-D**), a ribonucleotide reductase inhibitor, did not deplete CCDC137, even at the highest concentrations tested. Olaparib, a PARP1/2 inhibitor that prevents DNA damage repair and leads to an accumulation of DSBs, degraded CCDC137 only at high concentrations (**Figure S3E-F**). Among conditions that induced CCDC137 degradation, degradation inversely correlated with amount of DNA damage. However, CCDC137 degradation did not correlate with the absolute amount of damage present (**Figure 4E**). For example, in instances where camptothecin and cisplatin both induced higher γH2AX levels than Vpr, only camptothecin resulted in CCDC137 degradation. Degradation was also independent of the ability to cause G2/M arrest (**Figure 4F**) – consistent with the degradation of CCDC137 by Vpr mutants lacking this ability (**Figure 1**) – or activate p53 (**Figure 4G-H**). Together, these results suggest that CCDC137 depletion is a consequence of specific genomic insults.

**Figure 4.**
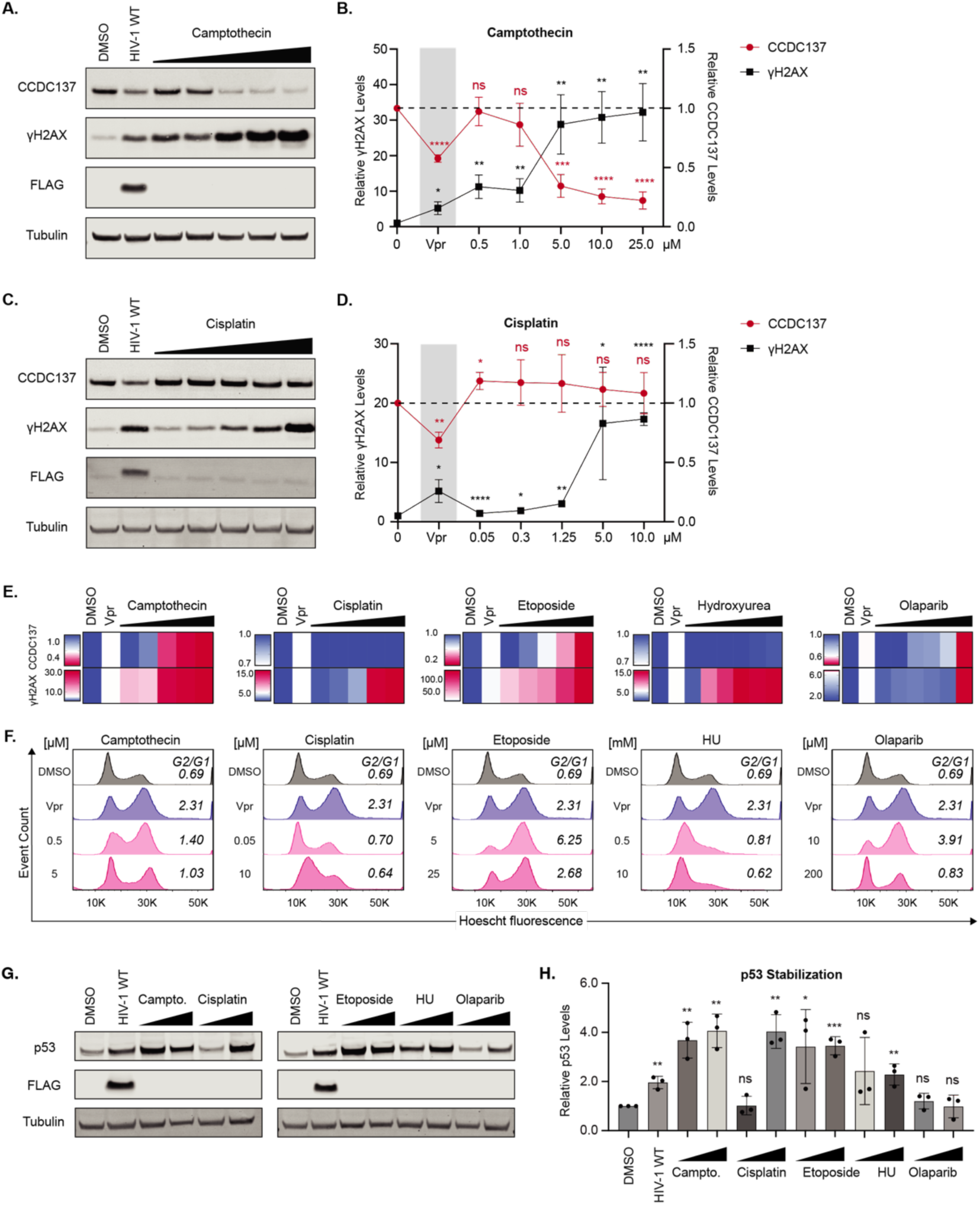
CCDC137 depletion is a consequence of specific types of DNA damage. (**A**) Representative western blot of CCDC137 depletion following 24hr of treatment with DMSO, HIV-1 WT, or titrating amounts camptothecin performed and (**B**) quantified as described in Figure 1C. (*n=3*) (**C**) Representative western blot and (**D**) quantification of CCDC137 and γH2AX levels in the presence of titrating amounts of cisplatin (*n=3*). (**E**) Heatmaps indicating CCDC137 and γH2AX levels in U2OS cells treated with DMSO, HIV-1 WT Vpr, or titrating amounts of the indicated genotoxic agents. Values for CCDC137 or γH2AX in a given condition are averaged from three separate experiments (*n=3*). Top row: color gradients depict the percentage of CCDC137 remaining in the empty vector control (dark blue), the lowest percent of CCDC137 measured among the represented conditions (dark pink), and CCDC137 levels in Vpr-treated cells as a midpoint (white). Bottom row: color gradients depict γH2AX induced by empty vector (dark blue), the highest γH2AX value measured among the given conditions (dark pink) and γH2AX levels induced by Vpr as a midpoint (white). (**F**) Representative flow cytometry plots of cell cycle profiles under the indicated conditions as described in Figure 1A. (**G**) Representative western blots and (**H**) quantification of p53 levels following 24hr of treatment with DMSO, HIV-1 WT, camptothecin (0.5µM or 5µM), cisplatin (0.05µM or 10µM), etoposide (5µM or 25µM), hydroxyurea (0.5µM or 10mM), or Olaparib (10µM or 200µM) (*n=3*). Results were analyzed by unpaired t-tests. Asterisks indicate statistical significance from empty vector control; error bars represent ± standard deviation (ns = not significant, *P≤0.0332, **P≤0.0021, *** P≤0.0002, ****P<0.0001). Campto. = camptothecin; HU = hydroxyurea.

### Vpr orthologs that degrade CCDC137 cause nucleolar stress

Considering that CCDC137 is a nucleolar protein, and there is extensive crosstalk between the DDR and nucleolus, we hypothesized that CCDC137 degradation may derive from DNA damage that specifically affects nucleolar pathways. In support of nucleolar involvement, of the genotoxic agents we assayed, only those that degrade CCDC137 – camptothecin, etoposide, and olaparib – perturb the nucleolus (37, 58–62). Cisplatin, however, acts via nucleolar-independent DDR activation (63), in line with its inability to degrade CCDC137. To test this hypothesis, we first asked whether Vpr causes nucleolar stress.

Nucleolar stress is diagnosed based on alterations to the nucleolar proteome and changes in nucleolar morphology and function (27). To assess the effects of Vpr on the nucleolar proteome, we leveraged two pre-existing studies that specifically assayed proteomic changes induced by Vpr (47, 48). Johnson *et al.* infected Jurkat E6.1 T cells with HIV-1 ΔEnv NL4-3 (WT), HIV-1 lacking one of each accessory gene Vpr, Vpu or Vif (ΔVpr, ΔVpu or ΔVif, respectively), or uninfected control cells and quantified changes in the total proteome. Gene set enrichment analyses (GSEA) of protein groups differentially increased or decreased among the different infection conditions revealed that the nucleolus and nucleolar processes harbored some of the most significantly decreased protein changes (**Figure 5A and Figure S4A**). When compared to mock infection, these changes were only lost with ΔVpr infection. When compared to WT infection, only ΔVpr, but not ΔVif or ΔVpu, were statistically significant for the decrease in proteins associated with nucleolus and nucleolar processes, together showing Vpr is necessary and sufficient for alterations to the nucleolar proteome.

**Figure 5.**
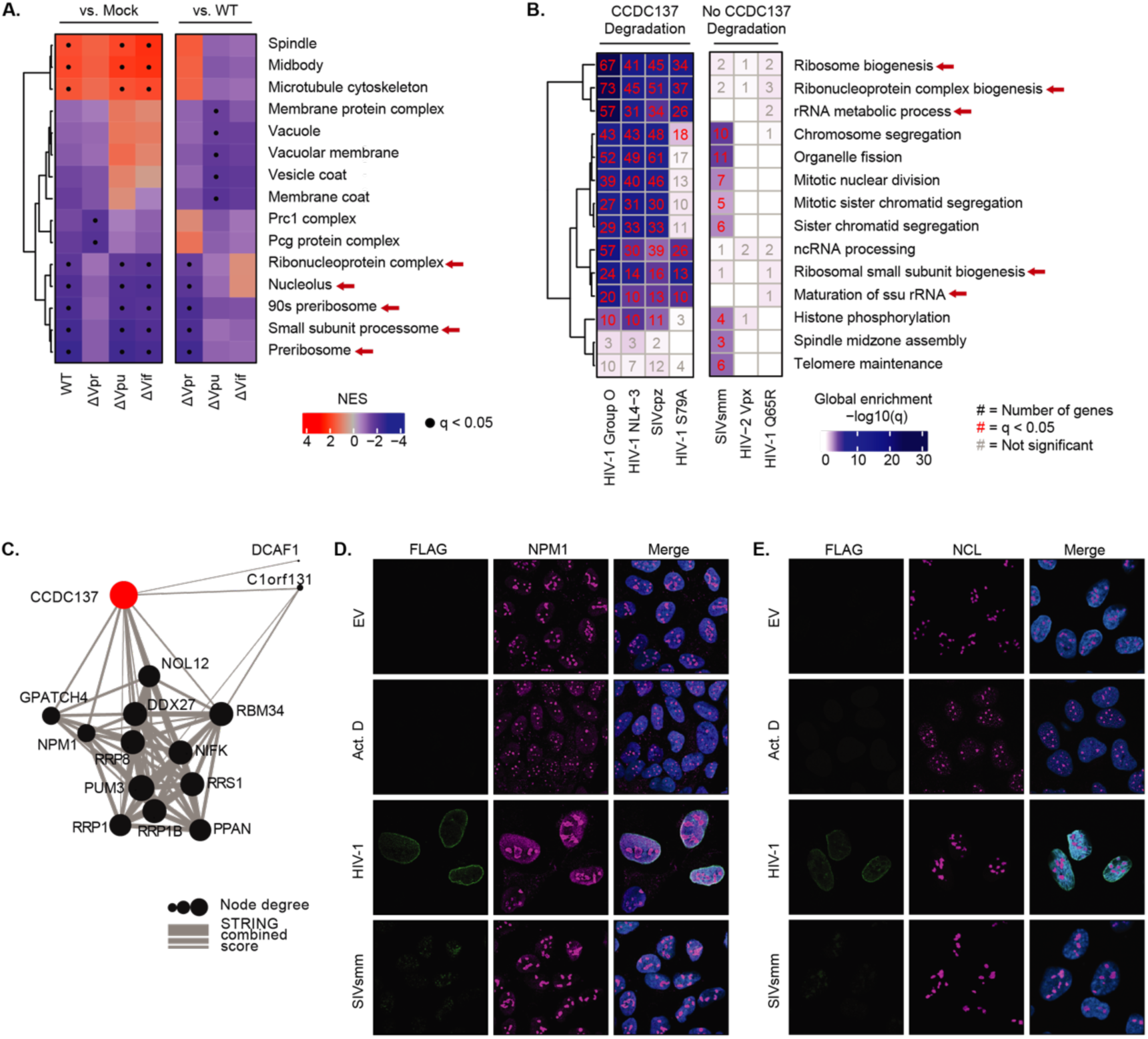
Vpr induced nucleolar stress correlates with CCDC137 degradation. (**A**) Gene set enrichment analysis (GSEA) of all expressed genes in (48) abundance proteomics dataset, using gene sets from the GO Cellular Component ontology. Graph legend shows normalized enrichment score (NES) computed using the enrichment and adjusted p-value values. Red arrows highlight nucleolar-associated gene sets. (**B**) GO terms enriched by gene set overrepresentation analysis (GSOA) (adjusted p-value < 0.05) of differentially expressed genes from (47) (log2FC > 0.5, Vpr expression versus empty). Top terms extracted from the GO Biological Process ontology. (**C**) A subnetwork extracted from STRING of CCDC137 interactions possessing a combined score greater than 0.4. (**D**) Representative immunofluorescence images of U2OS cells treated with the indicated conditions probed for NPM1 (magenta) or (**E**) NCL (magenta), 3XFLAG (Vpr) (green), and DAPI (blue). Images were taken at 63X magnification.

To understand whether Vpr-driven changes in nucleolar pathways correlated with CCDC137 degradation, we next analyzed proteomic data from Greenwood *et al.*, in which CEM-T4 T cells were transduced with Vpr proteins that could or could not degrade CCDC137. We performed gene set overrepresentation analysis (GSOR) of differential protein abundance in each condition. In the presence of Vpr proteins that degrade CCDC137, including HIV-1 Group O, HIV-1 NL4-3, the cell cycle deficient mutant HIV-1 S79A, and SIVcpz Vpr, the most significantly downregulated cellular pathways were nucleolar processes (**Figure 5B**), particularly ribosome biogenesis. Similarly, cellular components associated with nucleolar functions harbored the most significantly depleted genes (**Figure S4B**). In contrast, among changes induced by Vpr that did not degrade CCDC137 – SIVsmm Vpr, the functionally dead mutant HIV-1 Q65R Vpr, and HIV-2 Vpx – neither nucleolar pathways nor cellular compartments harboring nucleolar processes were significantly downregulated. Further, nucleolar proteins make up the majority of proteins significantly altered by CCDC137-degrading Vpr in both the Johnson *et al*. and Greenwood *et al.* datasets (**Figure S4C**).

To determine whether CCDC137 interacted with nucleolar genes identified in the Greenwood *et al.* abundance dataset, we performed a functional protein association network analysis using STRING (51) (**Figure 5C**). We discovered that CCDC137 interacted with multiple nucleolar proteins, including those with known roles in rRNA processing and HIV infection (65–68). One of these proteins, nucleophomism (NPM1), is a prominent nucleolar stress sensor (64). Together, these analyses suggest that nucleolar disruption is a specific function of Vpr, and one unique to Vpr proteins capable of CCDC137 degradation.

We next determined whether Vpr triggered functional changes in nucleolar morphology that are canonically associated with nucleolar stress, including redistribution of NPM1 and nucleolin (NCL) from the nucleolus to the nucleoplasm. We focused on HIV-1 and SIVsmm Vpr orthologs based on their contrasting effects on nucleolar pathways in **Figure 5B** and human CCDC137 degradation phenotypes (**Figure 3A-B**). As positive controls for nucleolar disruption, cells were treated with 5nM Actinomycin D (Act. D), which selectively inhibits rRNA transcription (69), and etoposide or camptothecin. Consistent with the ability to degrade CCDC137, HIV-1 but not SIVsmm Vpr led to NPM1 translocation despite inducing similar γH2AX levels (**Figure S5A**), suggesting NPM1 dispersal correlates with CCDC137 degradation (**Figure 5D**). Moreover, this dispersal resembled the morphological changes induced by etoposide and camptothecin rather than Act. D (**Figure S5B-C**), suggesting Vpr-induced nucleolar stress is likely not a direct derivative of RNA Pol I inhibition. While Act. D also disrupted NCL morphology, none of the Vpr orthologs or genotoxic agents altered NCL localization (**Figure 5E**), further implying specificity to the nucleolar changes observed. Act. D was sufficient to induce CCDC137 depletion, however, indicating that direct nucleolar insults also result in CCDC137 degradation (**Figure S5D**). Overall, these data suggest that HIV Vpr causes nucleolar stress by altering the nucleolar proteome and inducing the most prominent hallmark of nucleolar stress (64), NPM1 translocation.

### Vpr induces CCDC137 depletion through ATR-dependent nucleolar stress

Thus far, our results suggest that specific types of DNA damage are sufficient to disrupt the nucleolus and deplete CCDC137. Consequently, we sought to clarify the pathways connecting these processes. Central to the nucleolar-DDR crosstalk are the DDR signaling kinases ATM and ATR, which can be activated in response to both direct and indirect nucleolar insults (70, 71). To determine whether activation of either kinase was required for CCDC137 depletion, we asked whether Vpr-mediated CCDC137 degradation was rescued by inhibition of ATM via KU-55933 (ATMi) or ATR via VE-821 (ATRi). While ATMi had no effect on CCDC137 degradation, ATRi fully rescued CCDC137 degradation induced by both HIV-1 and HIV-2 Vpr (**Figure 6A-B**). This rescue was only observed for Vpr, given that neither inhibitor rescued CCDC137 degradation induced by camptothecin or etoposide (**Figure S6A**). Overall, these results demonstrate that CCDC137 degradation requires Vpr-induced ATR activation.

**Figure 6.**
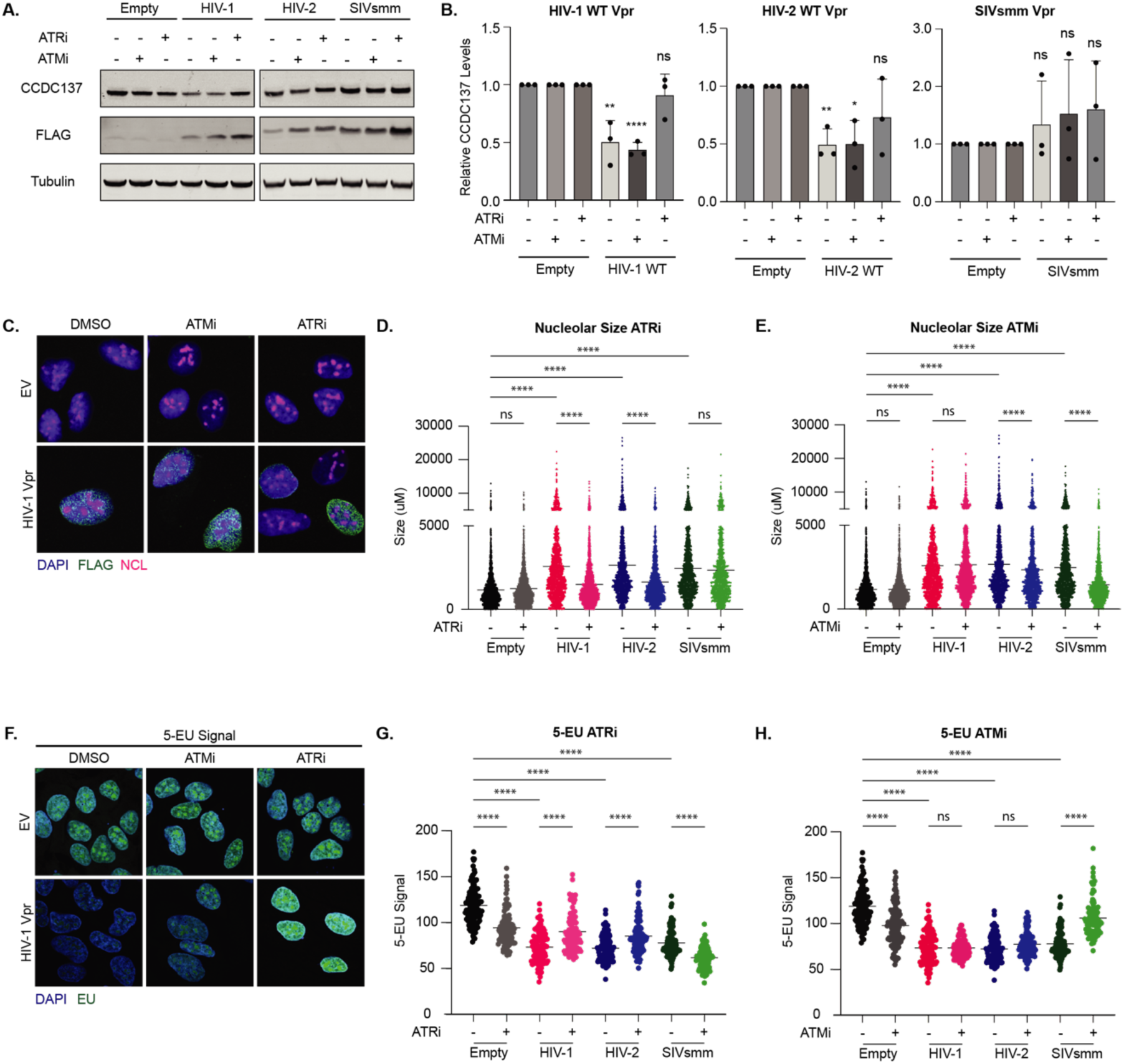
Vpr triggers ATR-dependent nucleolar stress thereby depleting CCDC137. (**A**) Representative western blots of CCDC137 levels in the presence of the indicated Vpr orthologs and DMSO, 10µM ATMi, or 10µM ATRi following a 24hr incubation. Western blots were probed and (**B**) quantified as in Figure 1 (*n=3*). Unpaired t-tests, error bars represent ± standard deviation, and asterisks indicate statistical significance from empty vector control; ns = not significant, *P≤0.0332, **P≤0.0021, *** P≤0.0002, ****P<0.0001. (**C**) Representative IF images of nucleolar size experiments probed for nucleolin (NCL) as described in Figure 5 (E). Images were taken at 100X. (**D-E**) Quantification of nucleolar size experiments performed as in (C) in ATRi (D) or ATMi (E) treated cells. Nucleolar area was measured for approximately 100 cells per condition. Representative quantification of two independent experiments (*n=2*). (**F**) Representative IF images of U2OS cells expressing HIV-1 WT Vpr or empty vector and stained for EU. Images were taken at 63X. (**G-H**) Quantification of total MFI in EU pulsed cells of the indicated conditions. MFI of EU signal was measured for approximately 100 cells per condition. Representative quantification of two independent experiments (*n=2*). Results in (D-E) and (G-H) were analyzed with a one-way ANOVA with Šidák’s multiple comparison test. Error bars represent ± standard deviation, and asterisks indicate statistical significance from empty vector control or cells subject to the same treatment conditions without inhibitor as indicated; ns = not significant, *P≤0.0332, **P≤0.0021, *** P≤0.0002, ****P<0.0001.

Given that ATR activation is necessary but not sufficient for CCDC137 degradation, we hypothesized that this degradation is due to the activation of an ATR-dependent nucleolar response to DNA damage. To test this hypothesis, we asked if Vpr-induced nucleolar stress was also dependent on ATR. We first looked at nucleolar size, which is indicative of nucleolar disruptions (27), including those caused by viral infections (26). Using the same Vpr and genotoxic agent panel as in **Figure 6A-B** and **S6A**, we detected larger nucleoli among all experimental conditions apart from Act. D, which decreased nucleolar area as expected (**Figure 6C-E and S6B-C**). Consistent with CCDC137 degradation, ATRi rescued the altered nucleolar sizes induced by both HIV-1 and HIV-2 Vpr, but not SIVsmm Vpr. Instead, the increased nucleolar size induced by SIVsmm CFU212 was fully rescued by ATMi, suggesting that Vpr orthologs that degrade CCDC137 do so by triggering a nucleolar ATR pathway. We also found that camptothecin augmented nucleolar size in an ATRi dependent manner, whereas etoposide induced changes were ATMi dependent (**Figure S6B-C**). This increased nucleolar size was not solely the result of enlarged nuclei, as nucleolar number also decreased, indicative of nucleolar fusion (**Figure S6D**). Moreover, it was not a consequence of G2/M arrest, as it was observed for SIVsmm CFU212 Vpr (**Figure 6D-E**), which does not induce arrest (**Figure S6E**).

Finally, we assayed whether Vpr disrupted ribosome biogenesis and whether this correlated with CCDC137 degradation. To determine this, we assayed changes in rRNA transcription, the rate limiting step of ribosome biogenesis (74), by 5-ethynyl uridine (EU) pulse-labeling. As this step can represent 80% of cellular transcription, general changes in EU signal are indicative of changes in nascent rRNA transcription (37). We found that HIV-1, HIV-2 and SIVsmm Vpr all significantly decreased rRNA transcription, thus disrupting the initial stage of ribosome biogenesis. Reflecting the nucleolar size assays, the decreased rRNA transcription induced by HIV-1 and HIV-2 Vpr was ATRi dependent whereas that induced by SIVsmm CFU212 was ATMi dependent (**Figure 6F-H**). Similarly, camptothecin and etoposide decreased EU signal, and these changes were ATRi and ATMi dependent, respectively (**Figure S6F-G**).

Together, these data show that Vpr orthologs that degrade CCDC137 cause nucleolar stress by modulating the nucleolar proteome, altering NPM1 localization and nucleolar size, and decreasing ribosome biogenesis. Moreover, they suggest that HIV Vpr degrades CCDC137 through the activation of a nucleolar-specific ATR pathway, suggesting CCDC137 plays a central role in nucleolar homeostasis.

## DISCUSSION

Vpr-mediated CCDC137 depletion represents a powerful example of how indirect host targets can reveal novel lentiviral accessory protein functions and help to define novel roles for cellular proteins. In this study, we establish CCDC137 as a non-canonical, but evolutionarily important, Vpr target that is part of a broader cellular response to nucleolar stress. By leveraging a set of HIV-1 Vpr mutants and genotoxic agents that do not cause G2/M arrest, we show that CCDC137 depletion is neither a direct cause or consequence of Vpr-induced cell cycle arrest. Further, we show that degradation is not mediated through canonical Vpr recruitment of the CRL4A^DCAF1^ ubiquitin ligase complex, and that Vpr and CCDC137 neither directly interact nor occupy the same nuclear compartments. By assaying degradation among Vpr and CCDC137 orthologs representative of the broader HIV and SIV diversity, we show that CCDC137 depletion is conserved among the HIV-1 but not HIV-2 lineage, suggesting it may constitute, or be a component of, an important function of Vpr from the pandemic HIV-1 lineage. CCDC137 depletion is also species-specific and CCDC137 harbors signatures of rapid evolution, consistent with a possible evolutionary arms-race. In defining how Vpr leads to degradation of CCDC137, we show that CCDC137 is depleted by Vpr orthologs and genotoxic agents that activate an ATR-mediated nucleolar stress response. We characterize CCDC137 depletion as one component of this nucleolar stress response and show that it occurs alongside other Vpr-induced nucleolar insults, including 1) modulated nucleolar pathways, 2) altered nucleolar morphology, and 3) downregulated rRNA transcription. When nucleolar stress is not accompanied by CCDC137 degradation, as in the case of SIVsmm Vpr, it is ATM dependent. Together, we identify a current model where Vpr activates a nucleolar-specific ATR signaling pathway that triggers nucleolar stress and, with it, CCDC137 degradation.

### Nucleolar stress may bridge Vpr-induced DNA damage and G2/M arrest

Here, we clarify that CCDC137 depletion is neither a direct cause nor consequence of cell cycle arrest. Yet, there is an apparent, albeit complex, relationship between these two phenotypes, and the temporal order of CCDC137 degradation, arrest, and nucleolar stress remains unresolved. Our results favor a process in which CCDC137 depletion precedes arrest: Vpr causes ATR-dependent nucleolar stress, of which CCDC137 degradation is a component, and this eventually triggers G2/M arrest. Given that CCDC137 depletion is not sufficient to trigger arrest, one possibility is that Vpr-induced nucleolar insults need to be severe enough to stall cell cycle progression. In support of this, the nucleolar stress response can activate an ATR-Chk1-mediated G2 arrest once a certain threshold of stress has been reached (75). The signatures of nucleolar stress we observe are not solely consequences of arrest, given that we see NPM1 redistribution and decreased rRNA transcription among conditions with varying arrest phenotypes. Moreover, through bioinformatic analyses, we also determine that downregulation of ribosome biogenesis and other nucleolar pathways is a specific cellular consequence of Vpr but not Vif, despite both proteins causing similar cell cycle arrest (76). In contrast to our model, others have shown that CCDC137 degradation can be caused by cell cycle arrest (52), though this may be due to the intertwined nature of arrest and nucleolar disruption, as ribosome biogenesis both regulates and is regulated by cell cycle progression. Such sensitivity of the nucleolus to disruptions may also explain why some Vpr orthologs are able to degrade only certain CCDC137 proteins in the same system. Overall, future work will be needed to establish a clearer temporal order of CCDC137 degradation, nucleolar disruption, and arrest.

By comparing the effects of Vpr and different genotoxic agents on CCDC137 levels and nucleolar integrity, our data also provide insight into how Vpr engages the DDR. Previous studies from our lab and others have shown that Vpr causes DNA damage and simultaneously activates and represses DDR pathways (16). Here, we have demonstrated that Vpr activates a nucleolar-specific ATR pathway that is distinct from ATR triggered by general DNA damage, as all of the genotoxic agents we tested activate classical ATR (77–80), but few disrupt the nucleolus. Yet, it remains to be seen whether activation of this pathway demonstrates Vpr-mediated ATR dysregulation or represents one of numerous ATR signaling branches engaged by Vpr. Moreover, it is also possible that the nucleolar ATR pathway is not activated by DNA damage but by independent nucleolar insults. What these insults may be and how they could differentially activate ATR remains to be identified. We further identify multiple phenotypes shared by HIV-1 Vpr and the Topo I inhibitor camptothecin, including robust CCDC137 degradation, and activation of ATR-dependent changes in nucleolar size and rRNA transcription. Past work has shown that Vpr induces double-stranded DNA breaks by structurally altering DNA in a manner that is Topo I and ATR dependent (81), supporting a mechanism in which Vpr might inhibit Topo I activity to activate ATR signaling. In addition, camptothecin has immediate effects on rRNA processing (59), corroborating a role for CCDC137 in this pathway, though it remains unclear whether which, if either, camptothecin mechanism Vpr phenocopies.

### CCDC137 may act as a sensor of nucleolar disruption with implications for cancer progression

CCDC137 has been classified as an oncogenic RNA binding protein, but its cellular functions and precise roles in cancer progression are only beginning to be understood. Our results reveal that CCDC137 is depleted as a consequence of ATR-dependent nucleolar stress. Identifying the exact nucleolar pathways in which CCDC137 participates is the next outstanding question. Considering that the nucleolus is a cellular stress sensor, it is possible that CCDC137 degradation, like NPM1 redistribution, represents an early step in this stress response. However, it is also possible CCDC137 degradation is a consequence of specific nucleolar pathways being disrupted. Our proteomic and STRING network analyses suggest that CCDC137 may be involved in rRNA processing, the inhibition of which has been identified as a stress-dependent regulatory event in the maintenance of nucleolar integrity (82). This would support a model in which CCDC137 acts as a nucleolar stress sensor, is depleted in response to specific ATR-dependent stress signaling, and effects a pause in rRNA processing that triggers the eventual downregulation of other ribosome biogenesis pathways. Given that unprocessed rRNA is stored in the nucleolus (82), this may explain why we see increased nucleolar size with Vpr and other agents that degrade CCDC137. This storage mechanism allows for the resolution of nucleolar stress and eventual return to normal function; if the nucleolus cannot coordinate this pause in rRNA processing with other steps of ribosome biogenesis, however, it can enter a state of irreversible stress that leads to cell cycle arrest and death (83). It is this disconnect in the ability to resume versus not recover homeostasis that may explain why some Vpr proteins are able to degrade CCDC137 without further downstream consequences, such as cell cycle arrest. Such a disconnect would also have important implications in cancer. For instance, if CCDC137 is acting as a sensor of nucleolar stress that dampens nucleolar processes when depleted, it is plausible that CCDC137 overexpression would prevent this ATR-dependent nucleolar inhibition. In line with this, ribosome biogenesis is hyperactivated in cancer and has been causally associated with malignant transformation and cancer progression (84). Although there is much exploration to be done in eliciting the exact pathways involved, this study supports that CCDC137 may be a potential therapeutic target whose depletion may slow oncogenesis. Moreover, it clarifies the DNA damage pathways that influence the nucleolus and recognizes various genotoxic agents that may be used to target CCDC137 and impede nucleolar processes.

### Vpr is one of many diverse viral proteins that may hijack the nucleolus to enhance viral replication

Finally, our findings raise the question of how Vpr-induced nucleolar stress might favor viral replication. Viral disruption of the nucleolus is a broad phenomenon, and numerous proteins from both plant and animal viruses localize to the nucleolus, redistribute nucleolar proteins for use in viral replication, and either enhance or inhibit ribosome biogenesis (27, 85). For instance, in herpes simplex virus infection (HSV1), nucleolar proteins localize to viral replication centers and co-localize with the viral ICP8 protein to enhance viral replication (86). Similarly, the infectious bronchitis coronavirus (IBV) N protein localizes to the nucleolus (87), and IBV results in nucleolar enlargement similar to that seen with Vpr (88).

In the context of HIV, the nucleolar localizations of both trans-activating proteins Tat and Rev are crucial to HIV-1 replication (89–91). Within this subnuclear compartment, Tat impairs rRNA processing (92) and remodels the nucleolar proteome (93), and Rev facilitates the export of un- and partially spliced viral RNA (94). How the effects of Vpr on the nucleolus coordinate with these functions is unknown, though previous studies connecting Vpr to translational impairment may provide insight into such an interplay. Vpr is reported to repress cellular translation via a mechanism that does not impede the translation of viral mRNAs (95), and in doing so may ready nucleolar processes for optimal Tat and Rev exploitation. In addition, NPM1 mediates the nucleolar retention of Tat and prevents its association with the super elongation complex (96), and it is conceivable that Vpr-induced NPM1 redistribution may increase Tat nucleoplasm localization to enhance transcriptional elongation. This is consistent with the proposed role of CCDC137 depletion in HIV-1 transcription (17), which may occur as part of the stress response that releases Tat. Future work will be necessary to dissect the how Vpr-induced nucleolar stress influences other HIV processes and impacts viral replication.

While there is still much to understand about the roles of CCDC137, Vpr, and the nucleolus in viral replication and cellular homeostasis, our data begin to unravel these complex connections. Moreover, we have characterized CCDC137 as an early sensor of nucleolar insults, and, in doing so, have opened new avenues for discovering further CCDC137 functions and pathways involved in the nucleolar stress response.

## DATA AVAILABILITY

The CCDC137 alignment data underlying this article are available in FigShare at https://figshare.com/ and can be accessed with https://doi.org/10.6084/m9.figshare.26841640 and https://doi.org/10.6084/m9.figshare.26841634.

## AUTHOR CONTRIBUTIONS

Karly Nisson: Conceptualization, Formal analysis, Investigation, Methodology, Project administration, Validation, Visualization, Writing – original draft, Writing – review & editing. Rishi S Patel: Formal analysis, Investigation, Writing – review & editing. Lucie Etienne: Conceptualization, Data curation, Formal analysis, Investigation, Resources, Writing – review & editing. Yennifer Delgado – Formal analysis, Investigation. Mehdi Bouhaddou – Formal analysis, Resources, Supervision. Oliver Fregoso: Conceptualization, Funding acquisition, Project administration, Supervision, Visualization, Writing – original draft, Writing – review & editing.

## ACKNOWLEDGEMENTS

We are grateful to Dr. Jefferey Long and Dr. Steve Jacobsen for use of the LSM980 and Dr. Peter Bradley for use of the Zeiss Axio Imager Z1. We thank Dr. Steven Bensinger for use of the LightCycler 480 system. We thank the James B. Pendleton Charitable Trust for providing the LiCOR Odyssey M. We thank Dr. Randilea Nichols Doyle and Vivian Yang for their comments on the manuscript. We thank Andrea Cimarelli and LP2L members (CIRI) for support and discussion. We thank Laurent Guéguen (LBBE) for continuous support maintaining the DGINN pipeline. We thank all the contributors of the publicly available bioinformatic programs and genomic sequences. The graphical abstract was created with Biorender.com.

