## Supplemental Figures for "HIV Vpr activates a nucleolar-specific ATR pathway to degrade the nucleolar stress sensor CCDC137"

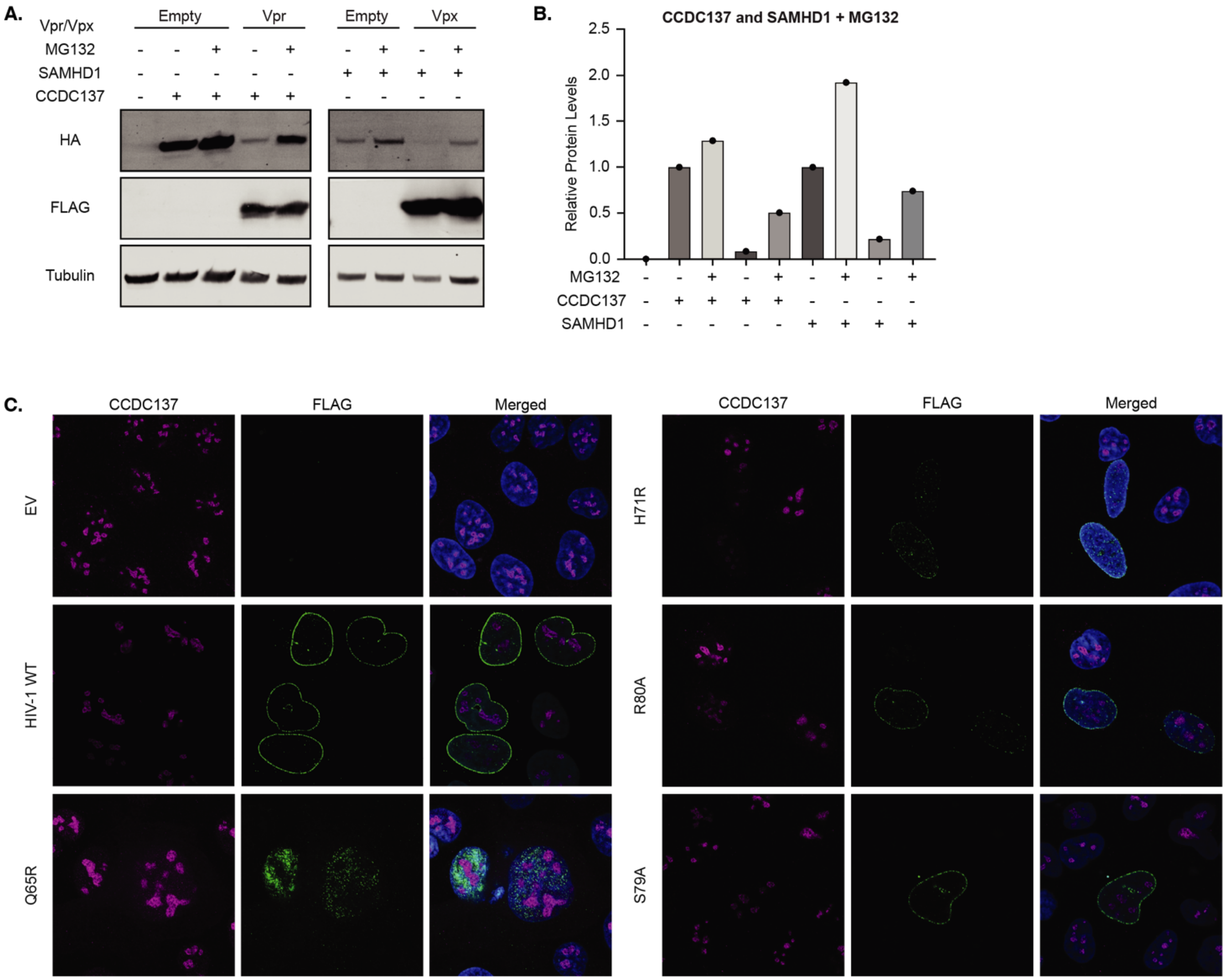


**Supplement Figure 1.**

(**A**) Western blot showing HA-CCDC137 or HA-SAMHD1 levels following transient co-transfection with HIV-1 Q23-17 Vpr or HIV-2 Rod9 Vpx, respectively, or empty vector control in the presence of 10uM MG132 or DMSO. (**B**) HA-CCDC137 and HA-SAMHD1 levels in (A) quantified and normalized as in Figure 1 (F). (**C**) IF images of U2OS cells infected with rAAV expressing the indicated Vpr mutants. Cells were permeabilized prior to fixation at 28hpi among all conditions except for the Q65R mutant, for which these steps were performed in reverse due to this mutant's exclusive presence in the soluble fraction. Cells were probed for endogenous CCDC137 (magenta), 3X-FLAG (green) and DAPI (blue) and imaged at 63X. Related to Figure 1.

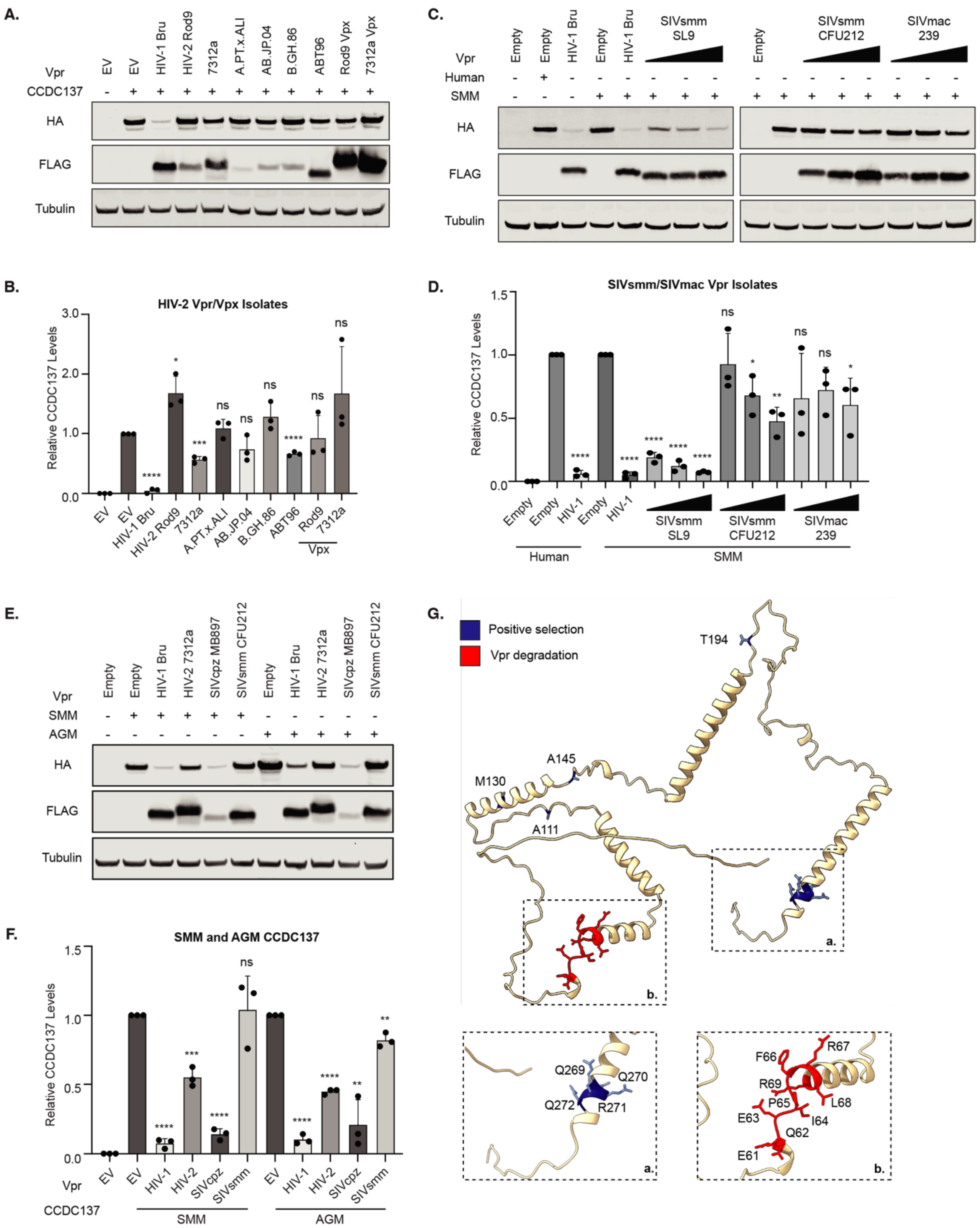


**Supplement Figure 2.**

(**A**) Representative western blots of human HA-CCDC137 degradation by HIV-2 Vpr isolates probed and (**B**) quantified as in Figure 2. (**C**) Representative western blots and (**D**) quantification of human or sooty mangabey HA-CCDC137 degradation by SIVsmm or SIVmac Vpr isolates. (**E**) Representative western blot and (**F**) quantification of sooty mangabey (SMM) and African Green monkey (AGM) HA-CCDC137 degradation by a panel of HIV-1 Bru, HIV-2 7312a, SIVcpz MB897 and SIVsmm CFU212 Vpr orthologs. (**G**) Alphafold predicted protein structure of CCDC137 labeled with PS marks (blue) and a Vpr binding site (red). Insets represent the variable C-terminus (a.) and a Vpr binding site (b.). All results were analyzed by unpaired t-tests. Error bars represent ± standard deviation, and asterisks indicate statistical significance from empty vector control; ns = not significant, *P≤0.0332, **P≤0.0021, *** P≤0.0002, ****P<0.0001. (A-D) Related to Figure 2. (E-G) Related to Figure 3.

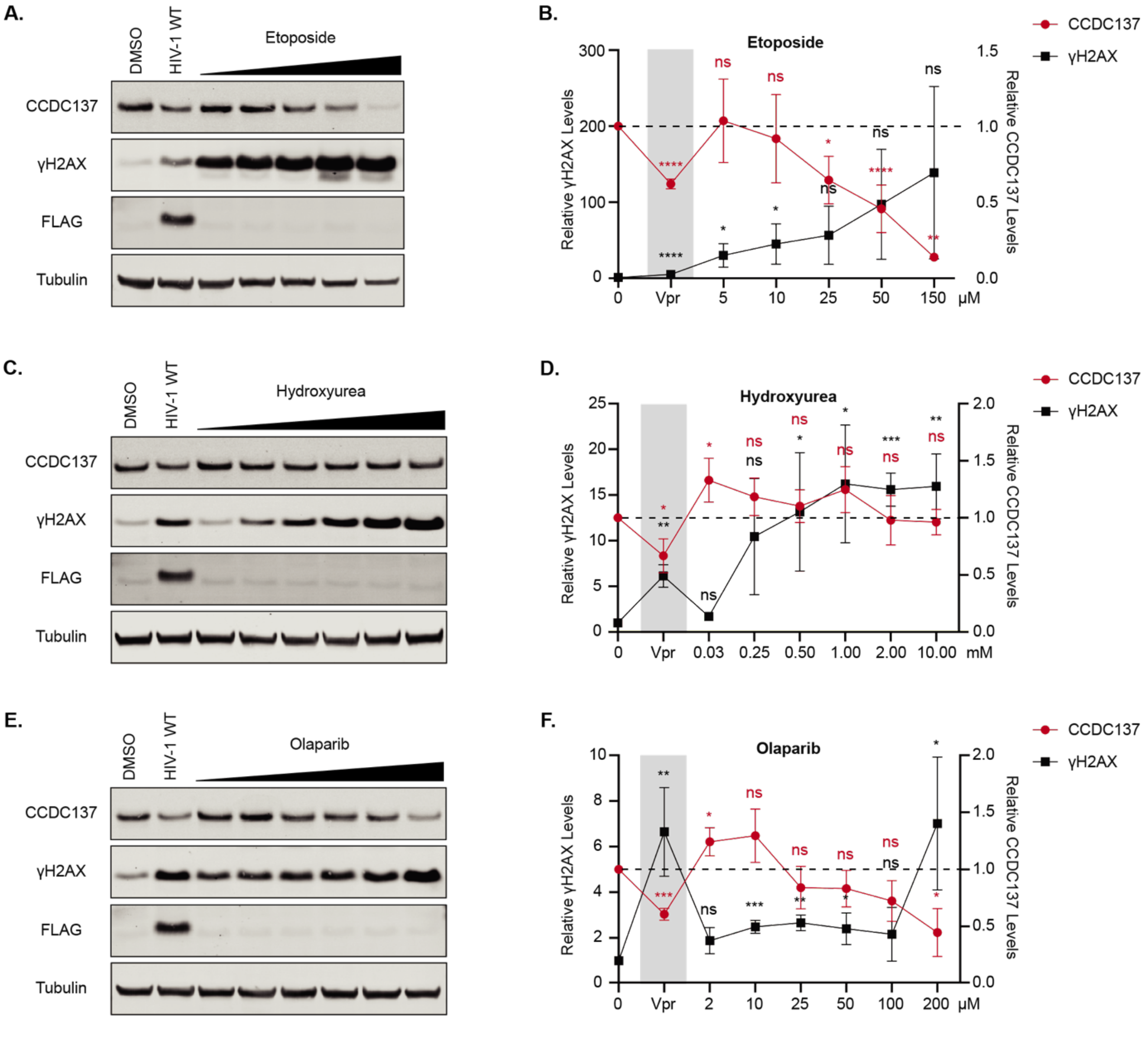


**Supplement Figure 3.**

Representative western blots and quantification of endogenous CCDC137 and γH2AX levels following 24hr of treatment with DMSO, HIV-1 WT Vpr or titrating amounts of etoposide (**A-B**), hydroxyurea (**C-D**), or olaparib (**E-F**). Western blots were probed and quantified as in Figure 4 (A-D). Quantification is representative of three separate experiments (*n=3*). All results were analyzed by unpaired t-tests. Error bars represent ± standard deviation, and asterisks indicate statistical significance from empty vector control; ns = not significant, *P≤0.0332, **P≤0.0021, *** P≤0.0002, ****P<0.0001.

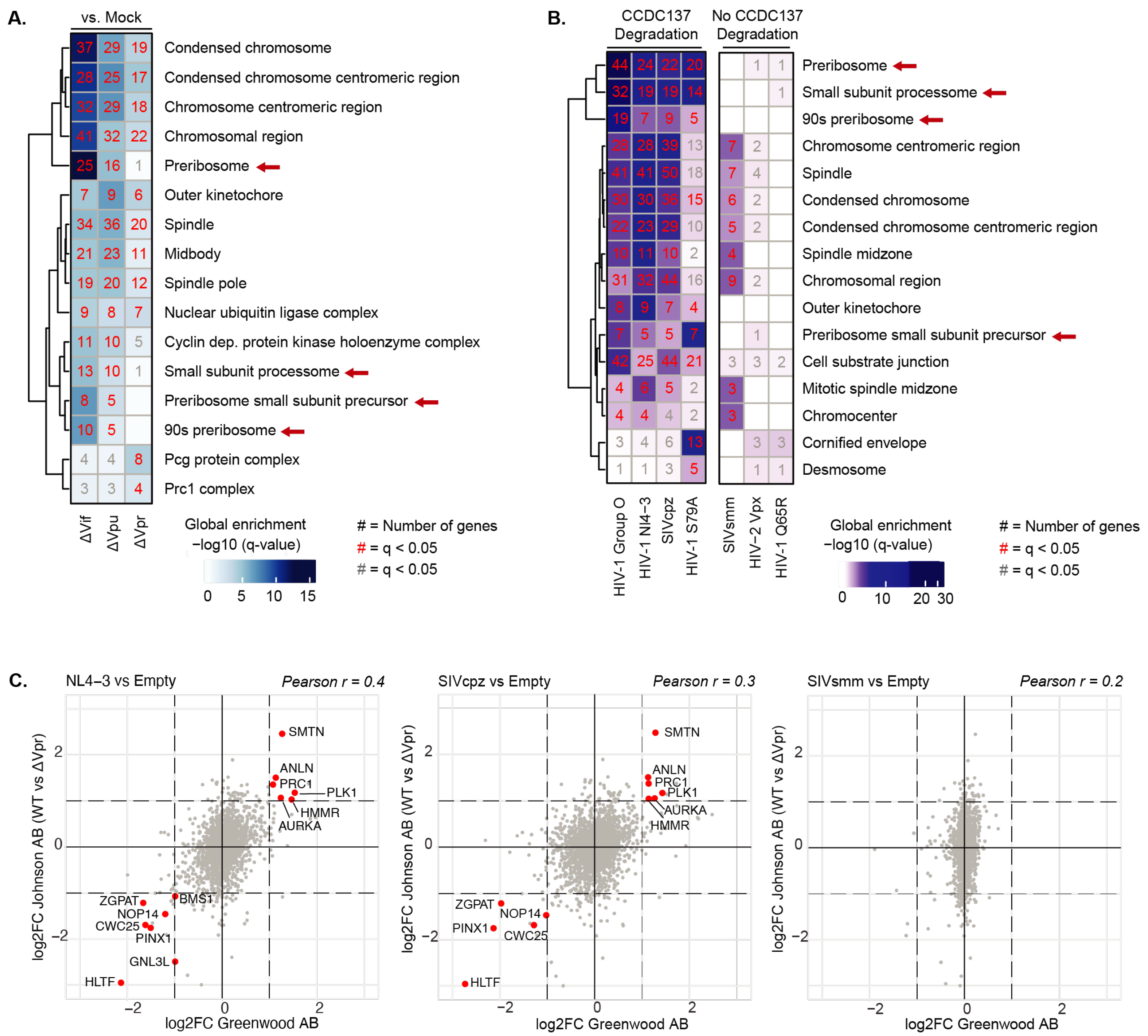


**Supplement Figure 4.**

(**A**) Enriched GO terms by GSOA (adjusted p-value < 0.05) of differentially expressed genes in the Johnson *et al.* dataset (log2FC > 0.5). Top terms represent Cellular Components. All numbers refer to the number of differentially expressed genes in a particular overrepresented GO term, and red numbers show gene sets with an adjusted p-value < 0.05. (**B**) Enriched GO terms by GSOA (adjusted p-value < 0.05) of differentially expressed genes in the Greenwood *et al.* dataset (log2FC > 0.5). Top terms represent Cellular Components. Numbers refer to the number of differentially expressed genes in a particular overrepresented GO term, and the red numbers show gene sets with an adjusted p-value < 0.05. (**C**) Up- or down-regulated proteins identified in both the Johnson *et al.* and Greenwood *et al.* abundance datasets. Jitter plots show comparisons between log2FC values from HIV-1 NL4-3 vs Empty, SIVcpz vs Empty, or SIVsmm vs Empty (X-axis), and the log2FC values from WT vs ΔVpr comparison (Y-axis). Red labeled dots represent genes that belong to the nucleolus term in the GO Cellular Component ontology and meet the following requirements: adjusted p-values ≤ 0.05, and log2FC>1 or log2FC<-1. Linear relationship between datasets was calculated using Pearson correlation coefficient. Related to Figure 5.

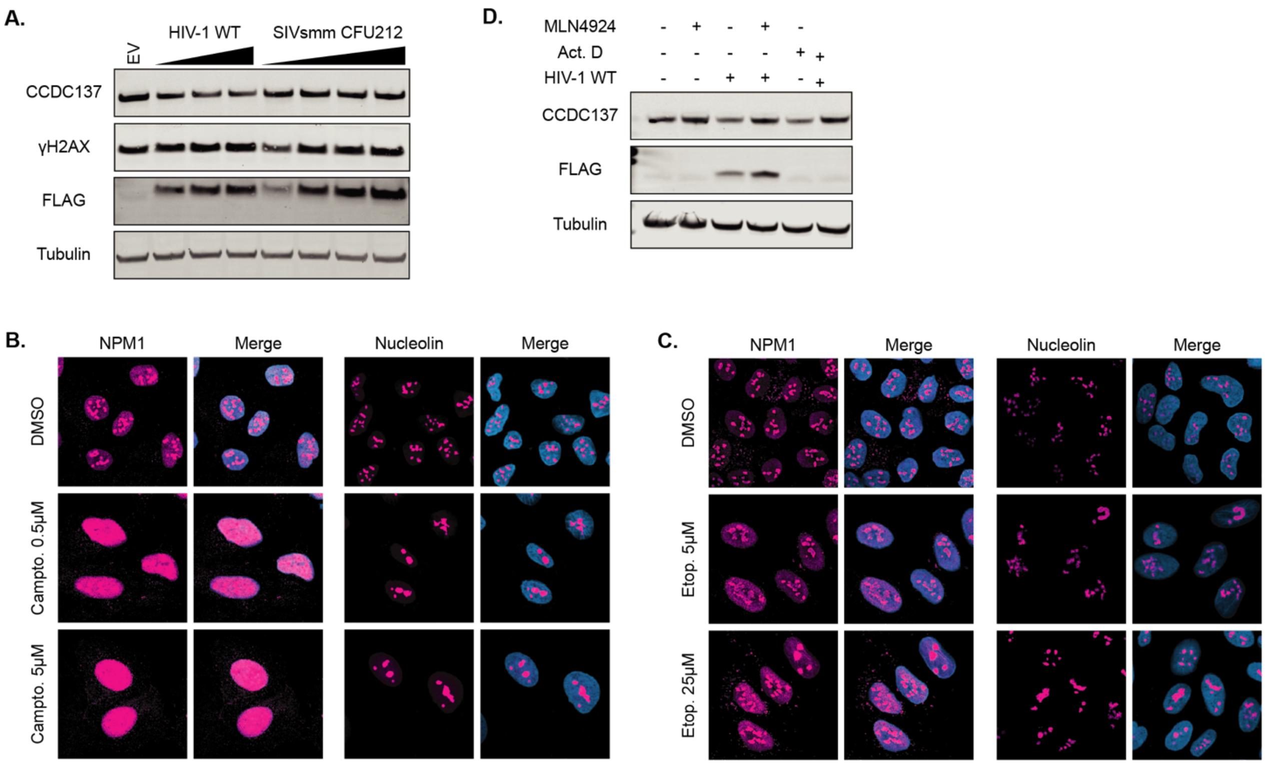


**Supplement Figure 5.**
(**A**) Western blot of endogenous CCDC137 and γH2AX levels induced by SIVsmm CFU212 and HIV-1 Q23-17 Vpr at 28hpi performed as in Figure 4. (**B**) Representative IF images of NPM1 and NCL localization in U2OS cells treated with 0.5µM or 5µM camptothecin and (**C)** 5µM or 25µM etoposide prepared as in Figure 5D. (**D**) Western blot of endogenous CCDC137 levels in cells treated with EV (- HIV-1 WT Vpr lanes) or HIV-1 Q23-17 Vpr in the presence of 5nM Actinomycin D (Act.D) and/or 10µM MLN4924. Related to Figure 5.

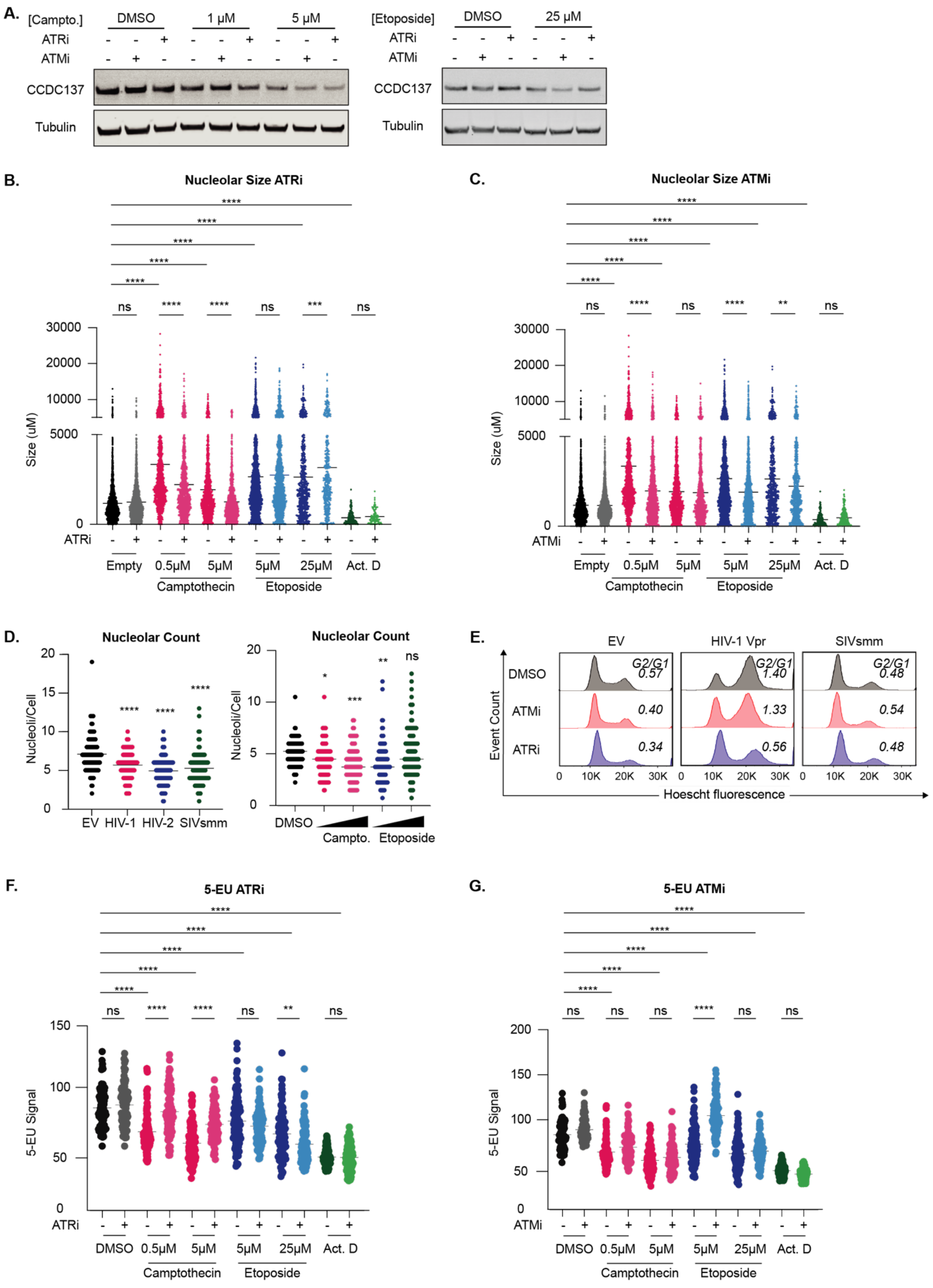


**Supplement Figure 6.**

(**A**) Western blots of endogenous CCDC137 levels in the presence of the indicated concentrations of camptothecin or etoposide incubated with DMSO, 10µM ATMi or 10µM ATRi following a 24hr incubation. (**B-C**) Quantification of nucleolar size experiments in ATRi (B) or ATMi (C) treated cells. U2OS cells were treated as (A) and nucleolar area was measured for 100-200 cells per condition. (**B**) Quantification of nucleolar number among U2OS cells incubated with the indicated Vpr or genotoxic agent as in Figure 6(D-E) and (B-C). The number of nucleoli/cell was measured for ~100 cells per condition. Results were analyzed with a one-way ANOVA with Dunnett's multiple comparison test. Error bars represent ± standard deviation, and asterisks indicate statistical significance from empty vector or DMSO control; ns = not significant, *P≤0.0332, **P≤0.0021, *** P≤0.0002, ****P<0.0001. (**C**) Representative flow cytometry plots of one cell cycle arrest experiment performed as in Figure 1A following 24hr incubation with the indicated Vpr and ATMi or ATRi. (**F-G**) Quantification of 5-EU experiments in ATRi (F) or ATMi (G) treated cells. Total MFI was measured for 100-200 cells per condition. Results were analyzed with a one-way ANOVA with Šidák's multiple comparison test. Error bars represent ± standard deviation, and asterisks indicate statistical significance from empty vector control or cells subject to the same treatment conditions without inhibitor; ns = not significant, *P≤0.0332, **P≤0.0021, *** P≤0.0002, ****P<0.0001. Related to Figure 6.

**File Supplement 1: Detailed primate CCDC137 evolutionary analysis.** Statistical results for the Detection of Genetic INNovations (DGINN) pipeline applied to 23 simian CCDC137 sequences. Bold values indicate tests that pass statistical thresholds for the indicated analysis. Bold sites are those that passed multiple analyses for significance of positive selection (PS), as indicated in "Summary of sites under PS" column. See Methods section for further details on the individual analyses performed. Related to Figure 3.
